# SMYD5 is a ribosomal methyltransferase which trimethylates RPL40 lysine 22 through recognition of a KXY motif

**DOI:** 10.1101/2024.10.10.616381

**Authors:** Joshua J. Hamey, Manan Shah, John D. Wade, Tara K. Bartolec, Richard E. H. Wettenhall, Kate G.R. Quinlan, Nicholas A. Williamson, Marc R. Wilkins

**Affiliations:** School of Biotechnology and Biomolecular Sciences, University of New South Wales, NSW, 2052, Australia; Florey Institute for Neuroscience and Mental Health, University of Melbourne, VIC, 3010, Australia; Melbourne Mass Spectrometry and Proteomics Facility, Bio21 Molecular Science and Biotechnology Institute, University of Melbourne, VIC, 3010, Australia; ARC Centre of Excellence for the Mathematical Analysis of Cellular Systems, University of New South Wales, NSW, 2052, Australia

**Keywords:** Ribosome, translation, lysine methylation, methyltransferase, substrate specificity

## Abstract

The eukaryotic ribosome is highly modified by protein methylation, yet many of the responsible methyltransferases remain unknown. Here we have identified SMYD5 as a ribosomal protein methyltransferase that catalyses trimethylation of RPL40/eL40 at lysine 22. Through a systematic mass spectrometry-based approach, we show that the human ribosome has 12 primary sites of protein methylation, including at RPL40 K22. Through *in vitro* methylation of synthetic RPL40 using fractionated lysate, we then identify SMYD5 as a candidate RPL40 K22 methyltransferase. We show that recombinant SMYD5 has robust activity towards RPL40 K22 *in vitro*, and that active site mutations ablate this activity. Knockouts of *SMYD5* in K562 cells show a complete loss of RPL40 K22 methylation and decrease polysome levels. By systematic analysis of its recognition motif, we show that SMYD5 requires a tyrosine in the +2 position, and thereby is incapable of methylating its previously reported histone substrates.

## Introduction

For over half a century, it has been known that the eukaryotic protein translational machinery is highly modified by protein methylation^1^. Only in the last two decades, however, have enzymes catalysing methylation of ribosomes and translation factors been discovered. As a result, it is now clear that the translational machinery is often a primary target of protein methylation systems in eukaryotes. In *Saccharomyces cerevisiae*, a suite of 17 protein methyltransferases are dedicated to modifying ribosomes and translation factors, usually generating stoichiometric methylation events^2^. In human, five methyltransferases targeting translation elongation factor 1A (eEF1A) have been discovered^3–9^, along with one translation elongation factor 2 (eEF2)-specific methyltransferase^10^ and an RPL3 histidine methyltransferase^11,12^. Additionally, RPL29 is a primary target of the lysine methyltransferase SET7/9^13^, and glutamine methylation of eukaryotic translation release factor 1 (eRF1) is a primary activity of HEMK2/N6AMT1/KMT9^14,15^. However, the full extent of protein methylation and cognate methyltransferases in the human translational machinery has yet to be mapped.

RPL40/eL40 is a late-joining protein of the 60S ribosomal subunit. RPL40 is encoded as a fusion gene with ubiquitin, which in humans is called UBA52. Cleavage of the fusion protein releases the N-terminal ubiquitin and results in the 52 amino acid RPL40 protein, one of the smallest ribosomal proteins, which joins the pre-60S subunit after its cytoplasmic export^16^. Nearly three decades ago, Williamson and co-workers found that rat RPL40/eL40 is trimethylated at lysine 22 (position 98 in the ubiquitin-fusion protein)^17^. Notably, RPL40 K22 was found to be completely trimethylated in rat liver, brain and thymus, suggesting an essential role for this modification^17^. This methylation event has since been observed in cryo-EM structures of human ribosomes^18^. However, the enzyme responsible for RPL40 K22 methylation has remained unknown.

SET and MYND domain-containing protein 5 (SMYD5) is a lysine methyltransferase reported to methylate histone H3K36 and H4K20^19,20^. However, deletion of SMYD5 results in only a small reduction in either of these modifications *in vivo*^19,21^, and SMYD5 shows weak or no activity towards histones *in vitro*^19,20,22^, especially when compared with canonical H3K36 and H4K20 methyltransferases SETD2 and SUV420H1^19^. This suggests that SMYD5 may not methylate these histone marks.

Here we have identified SMYD5 as the sole methyltransferase responsible for RPL40 K22 methylation in the human cell. We first identify RPL40 K22 methylation, along with 11 other methylation events, in actively translating ribosomes. Then, using biochemical fractionation of cell lysate in combination with *in vitro* methylation assays, we identify SMYD5 as a candidate RPL40 K22 methyltransferase. We then show that recombinant SMYD5 catalyses RPL40 K22 methylation *in vitro* and that knockouts of *SMYD5* result in a complete loss of RPL40 K22 methylation *in vivo*. In K562 cells, we show that *SMYD5* knockout results in a decrease in polysome levels, but does not decrease translation rate. By systematically profiling SMYD5 amino acid preference around RPL40 K22, we show that SMYD5 requires a tyrosine in the +2 position for substrate methylation, and that this explains its lack of activity towards histones H3 and H4 *in vitro*.

## Results

### Human ribosomes are predominantly methylated at 12 sites

To systematically profile protein methylation sites in the human ribosome, we employed heavy methyl SILAC (hmSILAC) labelling with ribosome isolation and a multi-protease LC-MS/MS approach (Figure 1A). K562 cells were cultured in the presence of light or heavy (^13^CD_3_–methyl) methionine, which we verified resulted in full labelling of a known methylpeptide from eEF1A (Figure S1). Light and heavy cells were mixed 1:1 and polysomes were isolated using Ribo Mega-SEC (Figure 1A, Figure S2). Ribosomal proteins from polysomes were then digested with trypsin, LysC, GluC or chymotrypsin, and peptides analysed by LC-MS/MS.

**Figure 1.**
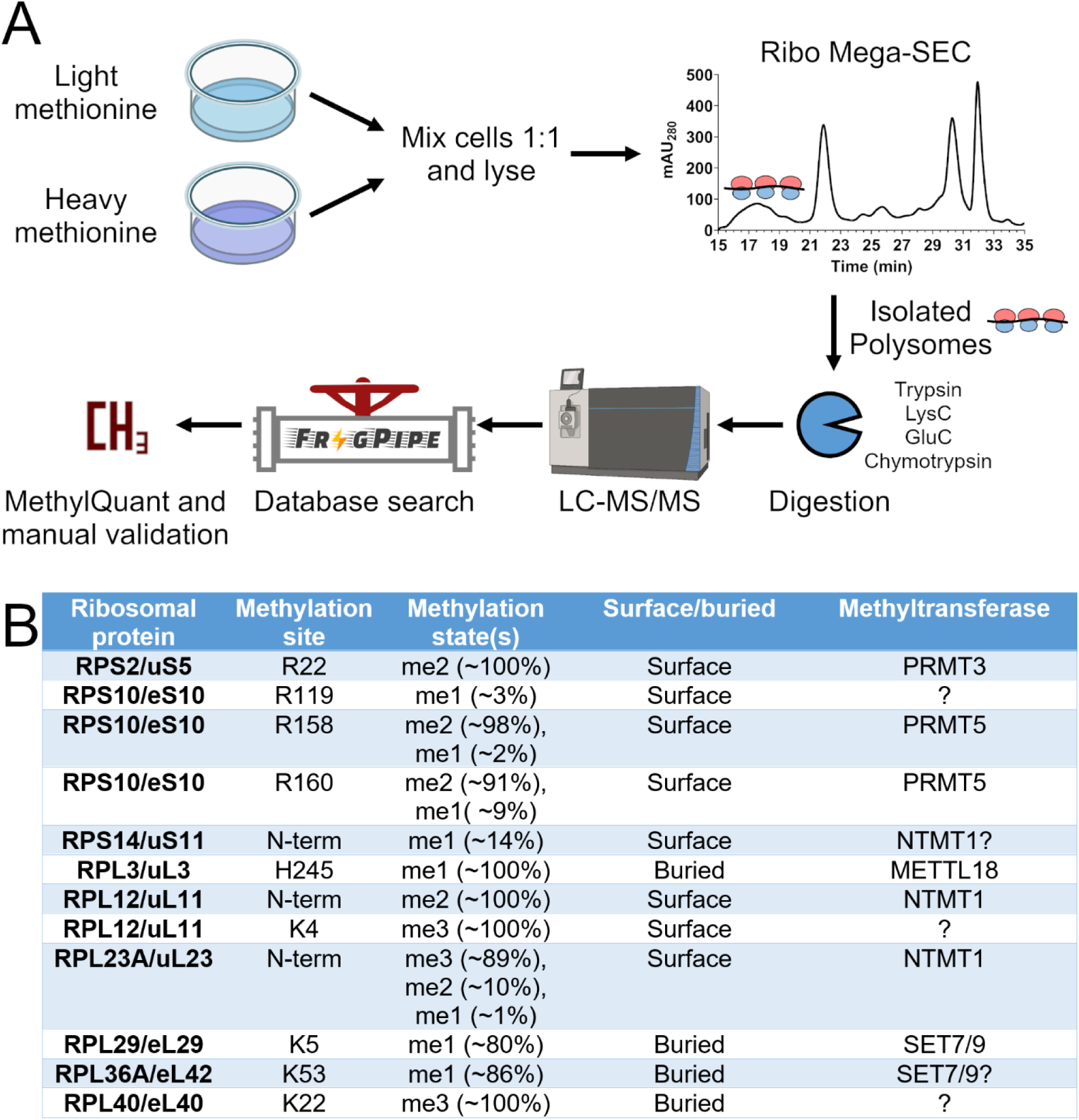
Systematic identification of human ribosome methylation sites. **A)** Workflow for analysis of protein methylation sites in the human ribosome. Cells were grown in the presence of light or heavy (^13^CD_3_-methyl) methionine, mixed 1:1 and lysed. Whole cell lysates were separated by Ribo Mega-SEC and the isolated polysomes were digested with four different proteases for analysis by LC-MS/MS. Putative methylation sites were identified by database searching, and *bona fide* methylation sites confirmed by the presence of light and heavy hmSILAC peptide pairs, as identified by MethylQuant and manual confirmation. **B)** Ribosome methylation sites identified in this study, their stoichiometries, surface-accessibility and upstream methyltransferase, if known.

Our analyses identified all 80 core ribosomal proteins with an average sequence coverage of ∼95% (Figure S3). From 130 ribosomal methylpeptide identifications, 81 were found to have genuine hmSILAC partners as determined using MethylQuant^23^ (Table S1). Spectra and heavy methyl SILAC pairs for representative methylpeptides are shown in Figures S4 and S5. Note that a putative series of lysine methylation events near the N-terminus of RPL23A were found to be entirely explained by N-terminal methylation, using targeted analyses with EThcD (Figure S4). We also further validated the identification of RPL40 K22 trimethylation through targeted analysis of a short tryptic peptide (sequence KCYAR) (Figure S6, Table S2). When results from all methylpeptides were summarised to non-redundant sites, 12 methylation sites were identified on 9 proteins, including four lysine sites, four arginine sites, three N-terminal sites and one histidine site (Figure 1B). The N-terminal methylation of RPS14 is novel and has not been previously reported. Almost all sites were of very high methylation stoichiometry, with the exception of RPS10 R119 and RPS14 N-terminal methylation (Figure 1B). We did not observe methylation of RPL4 K333 or RPS19 R67, as were reported in recent ribosome cryo-EM structures^24,25^, instead clearly observing two different unmodified peptides covering these residues in each case (Figure S7). This strongly suggests these sites are not methylated in human ribosomes, at least from K562 cells. Given the high sequence coverage in our data (Figure S3), the 12 sites reported here may be considered the canonical methylation sites of human ribosomes, along with RPS25 N-terminal methylation^26^.

### *SMYD5 methylates RPL40 at lysine 22* in vitro *and* in vivo

We sought to identify the methyltransferase responsible for the stoichiometric trimethylation of RPL40 K22. Full-length RPL40 was chemically synthesised and incubated with or without K562 lysate, in the presence of AdoMet, and any RPL40 K22 methylation was detected by mass spectrometry. A deuterated form of AdoMet (D_3_-methyl-AdoMet) was used in all assays to ensure the methylation mass shift was unique and distinguishable. Additionally, pooled non-ribosome-containing fractions from Ribo Mega-SEC was assayed against synthetic RPL40 in the same way. We detected methylation of RPL40 K22 in the presence of lysate or pooled SEC fractions, but not in their absence (Figure 2A, Figure S8). Next, we individually assayed the 17 non-ribosomal Ribo Mega-SEC fractions for activity against RPL40 (Figure S9). We detected strong methyltransferase activity in three of these fractions predominantly: F30, F31 and F32 (Figure 2B), suggesting that a single methyltransferase in the SEC fractions was methylating RPL40. We then quantified RPL40 K22 methylation generated by these fractions, as well as fractions F29 and F33, using parallel reaction monitoring (PRM) (Figure S10, top left), and also carried out deep proteomics analysis of SEC fractions F29 to F33. Across these five fractions, we detected a total of 4669 proteins, 4425 of which were quantified by label-free quantification (LFQ) (Table S3). Importantly, we detected 89 methyltransferase-like proteins in these fractions^27^, 85 of which were quantified by LFQ. We correlated the abundance of these 85 methyltransferases-like proteins to the PRM-quantified levels of RPL40 K22 methylation (Figures 2C, S10 and S11), and found that the top five highest correlating proteins were CIAPIN1, SMS, DPH5, SMYD5 and CDK5RAP1. Three of these, CIAPIN1, SMS and CDK5RAP1, are known to catalyse other reactions besides methylation^28–30^, and DPH5 is known to specifically methylate eEF2 in the formation of the diphthamide modification^31^. SMYD5 was therefore identified as the most likely RPL40 K22 methyltransferase from our data. In agreement with this, SMYD5 was previously identified as the strongest interactor of C-terminally tagged UBA52 (which, upon cleavage, would give C-terminally tagged RPL40) in a recent affinity purification-mass spectrometry screen^32^.

**Figure 2.**
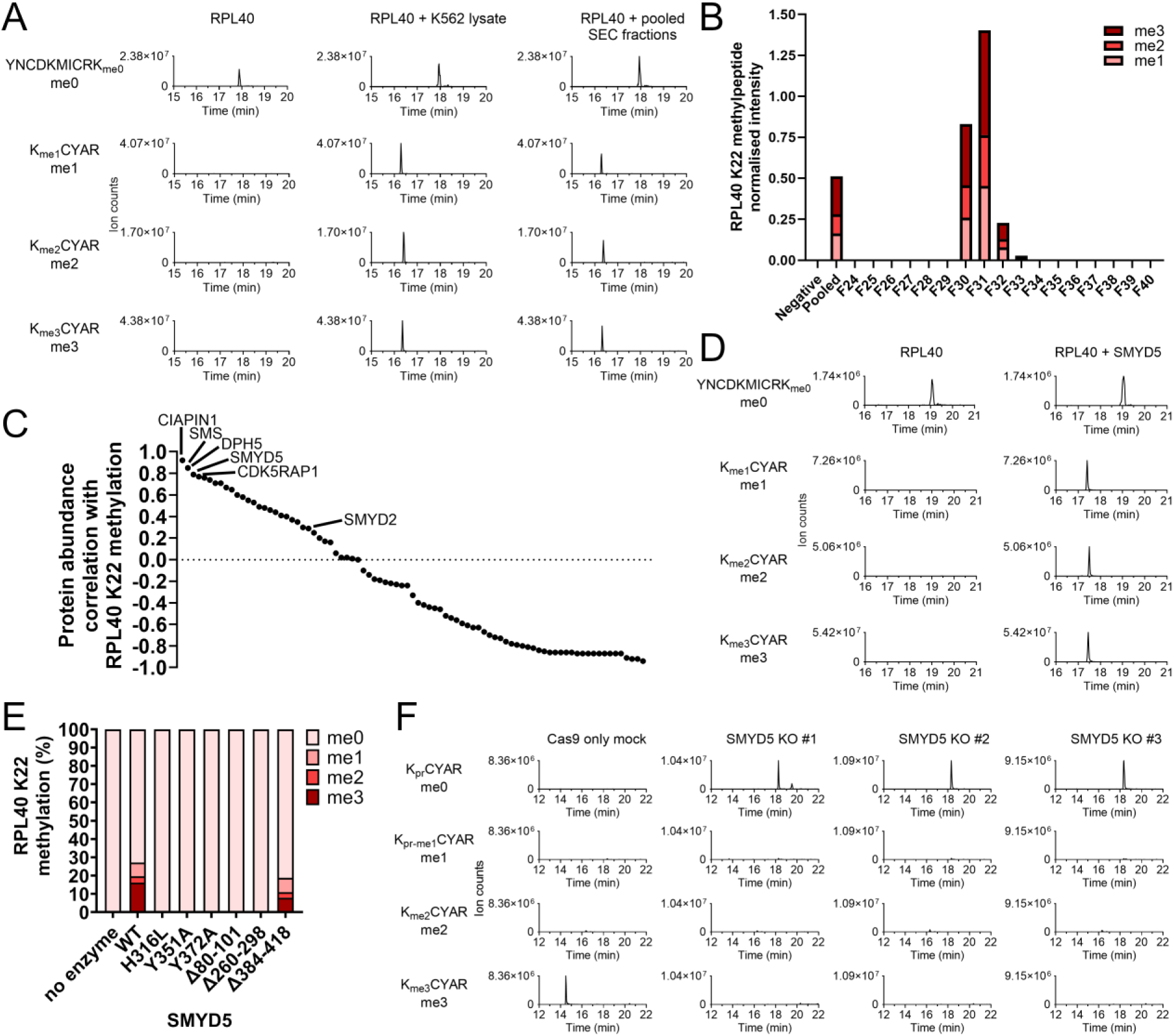
SMYD5 is the methyltransferase responsible for RPL40 K22 trimethylation. **A)** *In vitro* methylation assays of RPL40 by K562 lysate and pooled non-ribosomal Ribo Mega-SEC fractions. Synthetic RPL40 (200 μM) was incubated with K562 lysate (0.5 mg/mL) or pooled non-ribosomal Ribo Mega-SEC fractions (∼0.83 mg/mL) in the presence of D_3_-methyl-AdoMet. Proteins were separated by SDS-PAGE and the band corresponding to RPL40 was digested with trypsin and analysed by parallel reaction monitoring (PRM). Shown are the extracted ion chromatograms (XICs) for an unmethylated K22-containing peptide (sequence YNCDKMICR<u>K</u>, K22 underlined) and a CD_3_ mono-, di-, or tri-methylated K22-containing peptide (sequence <u>K</u>CYAR) (see Table S2). **B)** Methylation of RPL40 K22 by individual non-ribosomal Ribo Mega-SEC fractions (see Figure S9). Assays were carried out and RPL40 K22 methylation analysed as in (A). RPL40 K22 methylation levels in each assay were quantified by taking the area under the curve of XICs corresponding to the peptide <u>K</u>CYAR in its mono-, di- and tri-methylated state (all CD_3_-methyl), and by normalising to the abundance of the unmethylated peptide YNCDKMICRK. **C)** Pearson correlations between putative methyltransferase abundances and RPL40 K22 methylation levels, across fractions F29 to F33. See Figures S10 and S11 for all profiles. **D)** Recombinant SMYD5 catalyses RPL40 K22 methylation *in vitro*. Synthetic RPL40 was incubated with or without SMYD5 in the presence of D_3_-methyl-AdoMet for 2 h at 37 °C. RPL40 K22 methylation was then analysed as in (A). **E)** SMYD5 mutants (see Figure S12A) ablate its activity towards RPL40 K22. Recombinant SMYD5 mutants (see Figure S12B) were incubated with synthetic RPL40 in the presence of D_3_-methyl-AdoMet. RPL40 was propionylated and digested with trypsin, and the resultant levels of RPL40 K22 methylation were monitored by PRM of peptide KCYAR (see Table S2). The corresponding XICs are shown in Figure S12C. **F)** Knockout of *SMYD5* in K562 cells results in a complete loss of RPL40 K22 methylation. Lysates from Cas9 only mock and *SMYD5* KO cells were separated by SDS-PAGE and the band corresponding to RPL40 was propionylated and digested with trypsin. RPL40 K22 methylation levels were then monitored by PRM of peptide KCYAR in all its methylated states (see Table S2).

To verify if SMYD5 was the methyltransferase catalysing RPL40 methylation *in vitro*, we expressed and purified recombinant SMYD5 and assayed it against synthetic full-length RPL40, again using D_3_-methyl-AdoMet. In the absence of SMYD5, RPL40 K22 was unmodified, however in the presence of recombinant SMYD5, RPL40 K22 was substantially mono-, di- and tri-methylated (Figure 2D). Mutations of predicted SMYD5 active-site residues (H316L, Y351A and Y372A) abolished activity towards RPL40 K22 *in vitro* (Figures 2E and S12). Additionally, deletion of structural elements unique to SMYD5 among the SMYD enzymes^22^ (i.e. the “M-insertion” (residues 80-101) and “S-insertion” (residues 260-298)) also abolished its *in vitro* activity, while deletion of its unique poly-E tail did not significantly affect activity (Figures 2E and S12D). Together, our data show that SMYD5 catalyses trimethylation of RPL40 at K22 *in vitro*.

To verify if SMYD5 is responsible for RPL40 K22 methylation *in vivo*, we generated *SMYD5* knockout (KO) K562 cell lines using two different guide RNAs (Figure S13). Three different *SMYD5* KO clones were generated (one from guide 1, two from guide 2), all with homozygous mutations in *SMYD5* predicted to result in frameshifts and premature stop-codons (Figure S13). Using PRM, we monitored RPL40 K22 methylation levels in *SMYD5* KOs and in a Cas9 only mock control cell line. While the mock control cells showed the expected stoichiometric trimethylation of RPL40 K22, all three *SMYD5* KO cell lines showed a complete loss of K22 trimethylation and the corresponding gain of unmethylated K22, with no detectable mono- or di-methylation (Figure 2F). This confirms that SMYD5 catalyses RPL40 K22 methylation in cells, and indicates that SMYD5 is the sole enzyme responsible for this methylation event.

### Loss of RPL40 K22 methylation reduces polysome levels

RPL40 K22 lies at the interface of RPL40 and the 28S rRNA, where the methyl groups are predicted to make contacts with the phosphate backbone of 28S rRNA residues G1945 and C4412 (Figure 3A, B). We therefore tested whether *SMYD5* knockout, and therefore loss of RPL40 K22 methylation, results in changes to translation. Using Ribo Mega-SEC, we profiled the levels of polysomes and monosomes in *SMYD5* KO cells compared with Cas9 only mock control cells, noting that the monosomes co-elute with the 60S subunit on the column used here (Figure S2). *SMYD5* KO cells showed a clear decrease in polysomes, with a corresponding increase in the monosome/60S peak (Figure 3C, Figure S14). This corresponds to a ∼30% decrease in the polysome-to-monosome/60S ratio (Figure 3D). As RPL40 has been shown to be substoichiometric in polysomes^33^, we then quantified ribosomal protein levels in polysome and monosome/60S fractions, in control and *SMYD5* KO cells. We observed no significant change in the relative abundance of RPL40, or any other ribosomal protein, in intact ribosomes (polysomes or monosomes/60S subunits) upon *SMYD5* KO (Figure 3E and S15). This indicates that loss of RPL40 K22 methylation does not affect ribosomal protein stoichiometry. Finally, we tested whether loss of RPL40 K22 methylation affects translation rate, using puromycin labelling of nascent polypeptide chains^34^. Under the conditions used, we observed no change in translation rate for *SMYD5* KO cells compared to Cas9 only mock control cells (Figure 3F). Overall, our data indicate that loss of RPL40 K22 results in a moderate decrease in polysome levels, but has no impact on ribosomal protein stoichiometry or translation rate in K562 cells.

**Figure 3.**
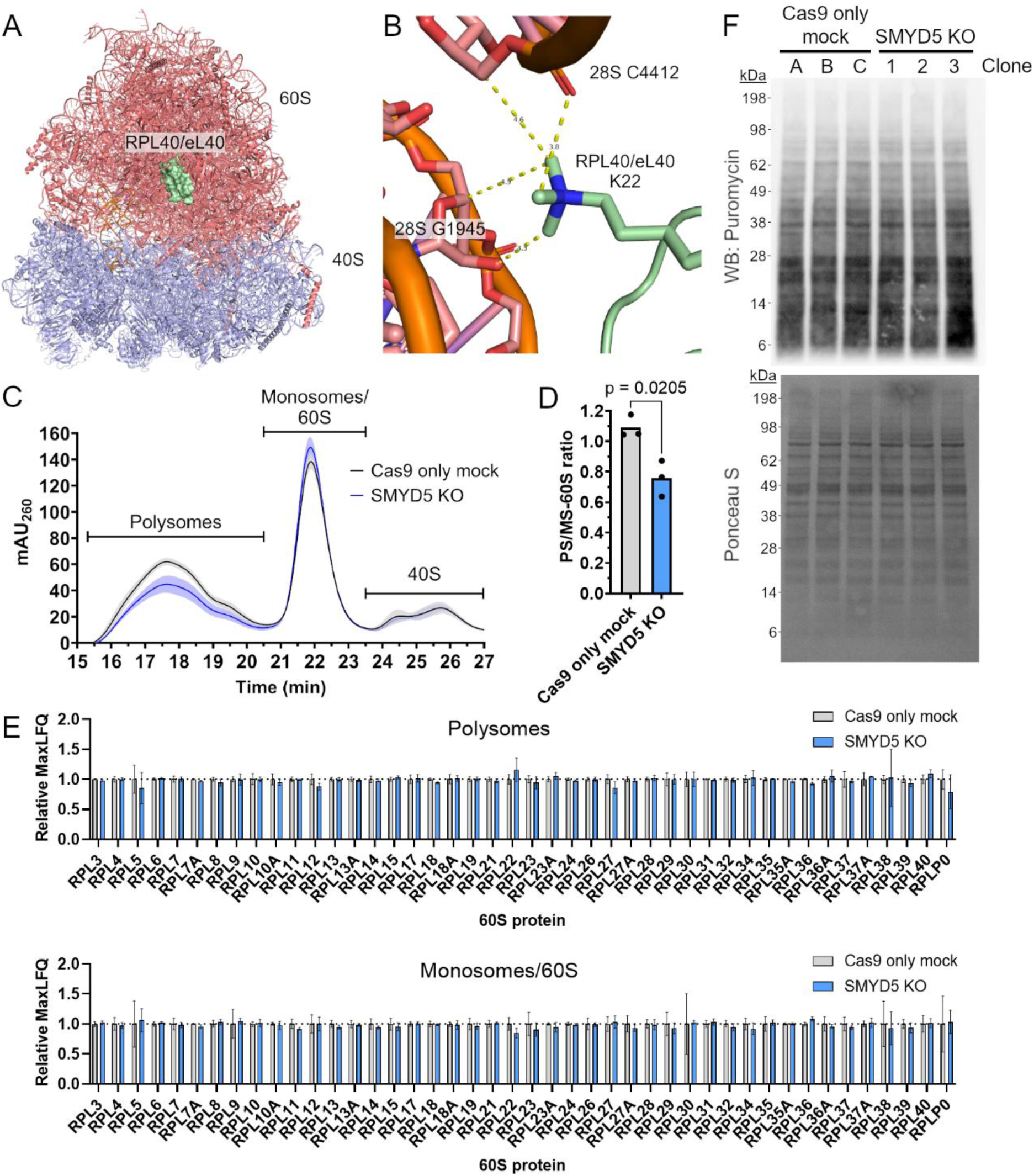
Loss of RPL40 K22 methylation decreases polysome levels, but does not affect translation rate or ribosomal protein stoichiometry. **A)** Ribosome structure (PDB ID: 8G5Y) showing location of RPL40 on the large subunit. **B)** Trimethylated RPL40 K22 makes contacts with the phosphate backbone of the 28S rRNA (PDB ID: 8G5Y). **C)** Averaged Ribo Mega-SEC profiles for Cas9 only mock controls (n = 3) and *SMYD5* knockout cell lines (n = 3), showing only ribosome fractions. Full profiles are shown in Figure S14. The shaded area represents one standard deviation. **D)** Ratios of the area under the curve of polysomes (PS) to monosome (MS)/60S peaks from Ribo Mega-SEC analyses, as shown in (C) and Figure S16. P-value is from a two-tailed t-test without equal variance. **E)** Polysomes or monosomes/60S fractions from Ribo Mega-SEC analyses in (C) were pooled and analysed by LC-MS/MS. MaxLFQ values for Cas9 only mock samples (n = 3) and *SMYD5* KO samples (n = 3) were normalised relative to the average MaxLFQ value for Cas9 only mock samples, separately for each ribosomal protein. Cas9 only mock and *SMYD5* KO samples were compared using unpaired t-tests with individual variances for each protein and multiple testing correction using the Holm-Šídák method. No significant differences were found (p > 0.05). Shown are 60S proteins; 40S proteins are shown in Figure S15. Error bars show one SD. **F)** *SMYD5* KO puromycylation assays. Three clones each of Cas9 only mock or *SMYD5* KO cells were incubated with puromycin for 15 min, and the resulting labelled nascent proteins were detected by immunoblotting against puromycin.

### SMYD5 does not methylate H3K36 or H4K20 as it recognises a KXY motif in its substrate

SMYD5 has been reported to methylate histone H3 at K36 and histone H4 at K20^19,20^. We therefore compared SMYD5 *in vitro* methyltransferase activity towards recombinant histones H3 and H4 against its activity towards RPL40. We again found that SMYD5 generated mono-, di- and tri-methylation of RPL40 at K22 (Figure 4A). In contrast, SMYD5 did not show any activity towards histone H3 at K36 or histone H4 at K20 (Figure 4A). We further assayed SMYD5 activity towards synthetic peptides corresponding to 15 residues around RPL40K22, H3K36 and H4K20, and found that SMYD5 robustly methylated the RPL40K22 peptide, but did not methylate either H3K36 or H4K20 peptides (Figure 4B, Figure S16). These data indicate that SMYD5 does not methylate H3K36 or H4K20 *in vitro*.

**Figure 4.**
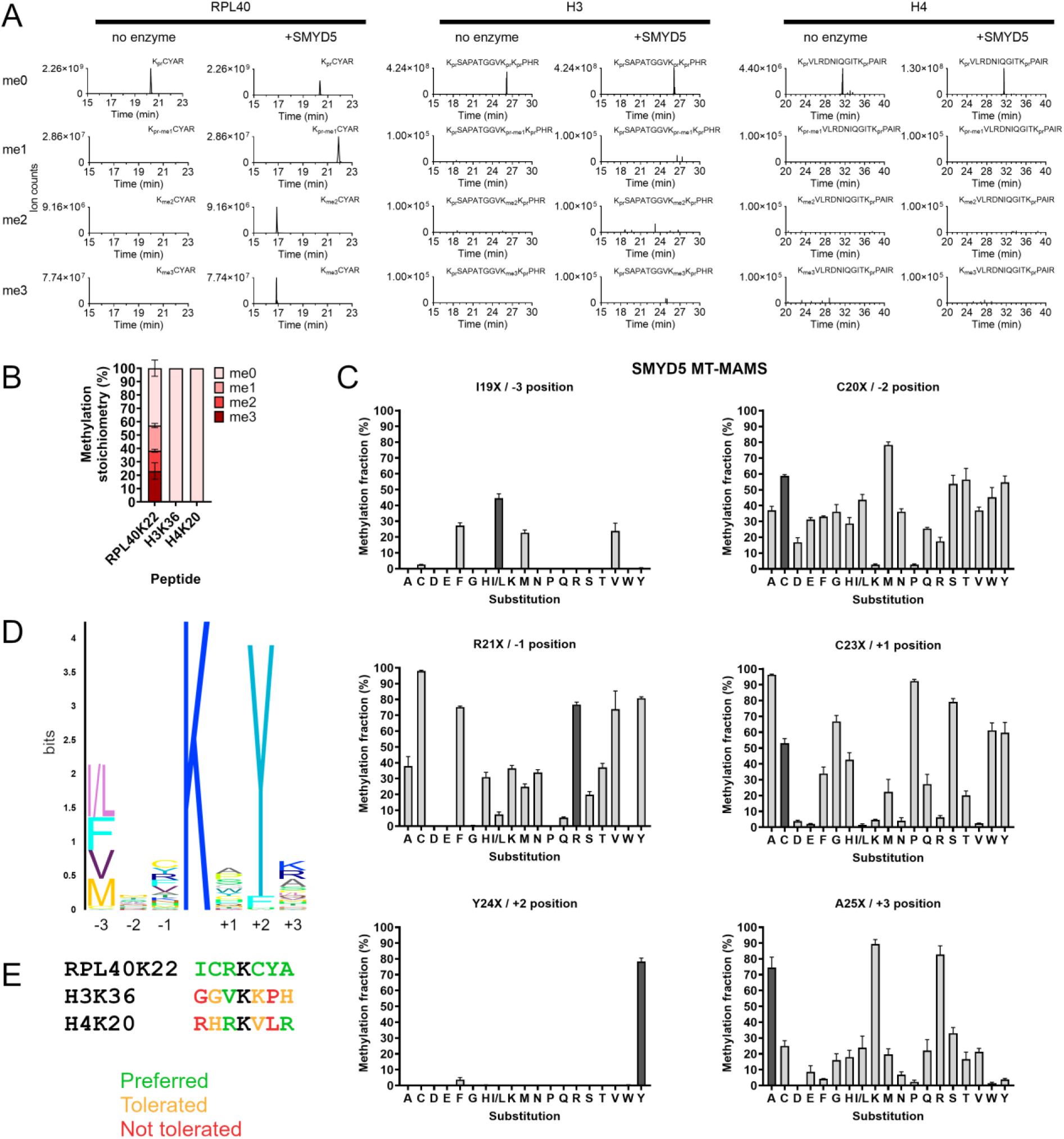
SMYD5 substrate recognition motif explains its lack of activity towards histone H3K36 and H4K20 *in vitro*. **A)** SMYD5 *in vitro* methylation assays of RPL40, histone H3 and histone H4. Synthetic RPL40, histone H3.1 or histone H4 were incubated with or without SMYD5 in the presence of D_3_-methyl-AdoMet. RPL40, H3 and H4 were subject to chemical propionylation and digested with trypsin. All methylation states of RPL40K22, H3K36 or H4K20 were then analysed using PRM targeting peptides KCYAR (RPL40K22), KSAPATGGVKKPHR (H3K36) and KVLRDNIQGITKPAIR (H4K20). **B)** SMYD5 *in vitro* methylation assays of peptides corresponding to RPL40K22, H3K36 and H4K20. Assays were carried out in triplicate and levels of SMYD5-catalysed methylation were quantified by LC-MS/MS. Error bars show one SD. **C)** SMYD5 Methyltransferase Motif Analysis by Mass Spectrometry (MT-MAMS). Peptide mixtures for each position (see Figure S17A) were methylated by SMYD5 *in vitro*, using D_3_-methyl-AdoMet, and methylation levels of all peptides were quantified by LC-MS/MS. Assays with SMYD5 were carried out in triplicate. Shown are the methylation fractions relative to 100% trimethylated RPL40K22; quantification of individual methylation states for all peptides is shown in Figure S17B. The amino acid found in the native RPL40 sequence is shown in dark grey. Error bars show one SD. **D)** Motif representation of SMYD5 amino acid preference. For each position, the total height of the stack represents the difference between the maximum possible entropy (log_2_ 19 ≈ 4.25) and the observed entropy, and the relative height of each residue is proportional to the relative methylation fraction of the peptide carrying that substitution compared to all peptides at that position. **E)** The local sequence context of H3K36 and H4K20 prevents their methylation by SMYD5. Preferred amino acids, shown in green, are those that resulted in a methylation fraction ≥50% of the most highly methylated peptide for that position; tolerated amino acids, shown in orange, are those that resulted in a methylation fraction <50%, but >0%, of the most highly methylated peptide for that position; not tolerated amino acids, shown in red, are those which resulted in a complete lack of methylation by SMYD5.

Given that SMYD5 methylates an RPL40 peptide, but not histone peptides, we reasoned that SMYD5 must recognise a linear sequence motif found around RPL40 K22 but not H3K36 or H4K20. To investigate this, we carried out MethylTransferase Motif Analysis by Mass Spectrometry (MT-MAMS)^35,36^ of SMYD5, using the RPL40 peptide as a template. Specifically, we profiled the effect of all single amino acid substitutions on SMYD5 activity, in three positions before and after K22, on the RPL40 peptide substrate (Figure 4C, Figure S17). Strikingly, we found that SMYD5 is highly selective for tyrosine at the +2 position, as found natively in RPL40, tolerating no other residue at this position except phenylalanine, which resulted in 20-fold less methylation (Figure 4C, D). We also found that SMYD5 is selective for large hydrophobic residues at the −3 position (Figure 4C, D), in particular leucine/isoleucine, in agreement with the isoleucine found at this position in RPL40. More generally, in each position we found that SMYD5 methylated the peptide containing the native RPL40 sequence at no less than 50% the level of the most methylated substituted peptide (Figure 4C). In contrast, H3K36 and H4K20 sequences contain many residues that SMYD5 does not prefer or does not tolerate, especially at the −3 and +2 positions (Figure 4E). More generally, when considering only ‘preferred’ residues at each position (i.e. residues allowing ≥50% methylation compared to the most methylated peptide for that position), SMYD5 recognises the following motif for positions −3 to +3: [FILMV]-[CILMSTWY]-[CFRVY]-<u>K</u>-[ACGPSWY]-Y-[AKR]. In the human proteome, only 19 proteins have this motif (Table S4), including RPL40, and none of the corresponding lysines have been detected as methylated in a range of methylproteome studies^37–42^. Overall, our data indicate that SMYD5 recognises, at a minimum, a KXY motif and thereby is unlikely to methylate histones as a preferred substrate.

## Discussion

The protein synthesis machinery is emerging as a primary target of protein methylation in eukaryotes. In agreement with this, we have here shown that SMYD5 is a ribosomal protein methyltransferase catalysing trimethylation of RPL40 K22. SMYD5 robustly catalyses mono-, di- and tri-methylation of RPL40 K22 *in vitro*, and knockouts of *SMYD5* result in a complete loss of RPL40 K22 methylation. In contrast to previous reports^19,20^, we detected no activity of SMYD5 towards histones H3 or H4, either as recombinant proteins or peptides.

Our results confirm two recent studies, which were published while this work was nearing completion^43,44^. Both groups also found that SMYD5 methylates RPL40 K22 but does not methylate histone substrates^43,44^. However, our discovery approach is complementary to the approaches taken in these studies, providing further confidence in the specificity of this enzyme-substrate relationship. Park et al. used *in vitro* methylation assays of SMYD5-depleted lysate to identify RPL40 as the sole target of this enzyme^43^, while Miao et al. screened a panel of SET domain-containing methyltransferases for activity towards recombinant RPL40^44^. In contrast, we used fractionated cell lysate as an enzyme source to *in vitro* methylate RPL40, and then proteomics to quantitatively correlate RPL40 K22 methylation levels with putative methyltransferase abundance across fractions. This provided an unbiased discovery of SMYD5 as the methyltransferase catalysing RPL40 methylation. Additionally, and in contrast to these two studies, our work provides insight into the mechanistic basis of SMYD5 preference for methylating RPL40 K22. Our systematic analysis of SMYD5 substrate recognition motif revealed that it requires a tyrosine in the +2 position for methylation, explaining its lack of activity towards H3K36 and H4K20, and suggesting a high degree of specificity for this enzyme.

We have shown that knockout of *SMYD5*, and the consequent loss of RPL40 K22, results in moderate (∼30%) decrease in polysomes, however we observed no change in translation rate. Given that monosomes are also known to be translationally active^45^, it is possible that an increased abundance of monosomes compensates for the decreased polysome levels. Our results partially agree with both Park et al. and Miao et al., who found that depletion of SMYD5 and the subsequent loss of RPL40 K22 methylation results in translational defects^43,44^. In contrast to our results, both these groups found that loss of RPL40 K22 methylation results in a decrease of translation rate^43,44^. In terms of the polysome profile, however, results from all three groups differ. While we showed a decrease in polysomes, Park et al. observed a slight increase in polysomes and a large decrease in 80S ribosomes, resulting in a significant increase in the polysome-monosome ratio upon loss of RPL40 K22 methylation^43^. In contrast, Miao et al. found that loss of RPL40 K22 methylation had no effect on 80S levels, but decreased the abundance of heavy polysomes and increased the abundance of light polysomes^44^. Most notably, they observed the presence of half-mer polysomes upon *SMYD5* knockout, which may indicate stalled pre-initiation complexes (PICs) due to deficient 60S subunits^44^. However, Park et al. found no defects in 60S biogenesis upon SMYD5 depletion^43^. Park et al. also found that loss of RPL40 K22 methylation results in slower elongation, as evidenced by a ∼30% increase in ribosome half-transit time^43^. This could be due to increased ribosome stalling, which agrees with the increased presence of disomes, indicative of ribosome collisions, observed by Miao et al.^44^. Overall, all studies show that RPL40 K22 methylation is critical for normal ribosome function, however there are differences in how this manifested, particularly in the polysome profile. It should be noted, however, that all three groups used different cell lines for these analyses.

SMYD5 adds to a growing number of protein methyltransferases that appear dedicated to methylating the ribosome and other translation-associated proteins^3–12^. Our SMYD5 MT-MAMS results show that the motif it recognises is very rare in the proteome, and in agreement with this, both Park et al. and Miao et al. found that *in vitro* assays with SMYD5 on SMYD5-depleted lysate produce a single radiographic band corresponding to RPL40^43,44^. It is therefore highly likely that SMYD5 is dedicated to methylating the ribosome, meaning there are now eight protein methyltransferases in human that are dedicated to methylating the translational machinery. As there are more ribosomal protein methyltransferases to be discovered (see below), it appears that, like in yeast^2^, the translational machinery is a primary target of the protein methylation system in human.

The fact that SMYD5 methylates the ribosome, rather than histones, prompts re-interpretation of some existing studies. For example, it has been shown SMYD5 is degraded during cold stress^46^ – while initially interpreted as a epigenetic regulatory mechanism, this may instead reflect the fact that translation elongation is reduced upon cold stress^47^. In another study, SMYD5 was shown to be associated with HIV-1 transcription^48^. However, it is known that viral replication is affected by translation^49^, and in particular, methylation of translation factors can affect viral replication^50^. SMYD5 has also been associated with embryonic stem cell differentiation, reported to be mediated through histone methylation^21,51^. However, the translational capacity of cells changes dramatically during differentiation, and differentiating cells are highly sensitive to perturbations of translation^52^. Given that SMYD5 methylation of RPL40 K22 can affect protein synthesis, its association with cold stress, HIV-1 transcription and differentiation may be secondary effects that result from changes to translational capacity in the cell.

Our systematic analysis of ribosomal protein methylation revealed 12 sites that are likely to constitute the canonical sites in the human ribosome, along with the known N-terminal methylation of RPS25^26^, for which we did not observe the corresponding N-terminal peptide. All these methylation sites were previously reported, except for N-terminal methylation of RPS14. Given that the RPS14 N-terminal sequence, A-P-R, contains the canonical [APS]-P-[KR] motif recognised by NTMT1 (also call NRMT1)^53^, it is highly likely that NTMT1 is responsible for this methylation site. Interestingly, in contrast with the near-stoichiometric trimethylation of the N-terminal A-P-K motif in RPL23A, RPS14 is subject to low stoichiometry monomethylation (Figure 1A), suggesting that A-P-R is a much less preferred motif than A-P-K *in vivo*. Therefore, there are four N-terminal methylation sites on human ribosomes, all likely catalysed by NTMT1. Of the four arginine methylation sites found here, RPS2 R22 is known to be catalysed exclusively by PRMT3^54^, while RPS10 R158 and R160 are methylated by PRMT5^55^. It is unclear which enzyme is responsible for RPS10 R119 methylation. RPL3 H245 methylation is catalysed by METTL18^11,12^ and RPL29 K5 methylation is catalysed by SET7/9 (SETD7)^13^. Although RPL36A K53 has also been reported to be methylated by SET7/9^56^, it is unclear if this enzyme is solely responsible for this methylation site, which notably does not sit within the known sequence recognition motif of SET7/9^57^. Therefore, with SMYD5 now known to catalyse RPL40 K22 methylation, there only remains three ribosome methylation sites without known or likely methyltransferases: RPS10 R119, RPL36A K53 and RPL12 K4.

## Acknowledgements

Mass spectrometry was performed at the Bioanalytical Mass Spectrometry Facility and fluorescence-activated cell sorting was performed at the Flow Cytometry Facility, both within the Mark Wainwright Analytical Centre of the University of New South Wales. M.R.W. acknowledges support from the Australian Research Council DP220102338.

## Author contributions

Conceptualisation, J.J.H., R.E.H.W., N.A.W., and M.R.W.; Methodology, J.J.H., M.S., and N.A.W.; Software, T.K.B.; Resources, J.D.W.; Formal Analysis, J.J.H. and M.S.; Investigation, J.J.H. and M.S.; Writing – Original Draft, J.J.H., M.S., and M.R.W; Writing – Review & Editing, J.J.H., M.S., J.D.W., T.K.B., R.E.H.W., K.G.R.Q., N.A.W., and M.R.W.; Visualisation, J.J.H. and M.S.; Supervision, K.G.R.Q. and M.R.W.; Funding Acquisition, M.R.W.

## Supplemental information

Document S1. Figures S1-S17 and Table S2.

Table S1. Methylpeptide identifications and MethylQuant analyses, related to Figure 1.

Table S3. Label-free quantification of proteins in Ribo-Mega SEC fractions F29-F33, and their correlation with RPL40 K22 methylation, related to Figure 2C.

Table S4. Matches to SMYD5 substrate recognition motif in the human proteome, related to Figure 4.

Table S5. Primers and DNA sequences used in this study.

## Data availability

Mass spectrometry proteomics data have been deposited to the ProteomeXchange Consortium via the PRIDE^58^ partner repository with the dataset identifier PXD056466. Target mass spectrometric data and Skyline files have been deposited to Panorama Public (https://panoramaweb.org/RPL40_methyltransferase.url) and ProteomeXchange (PXD056489).

## Methods

### Cell culture and heavy methyl SILAC labelling

K562 cells were maintained at 37°C with 5% CO2 and were cultured in RPMI-1640 supplemented with 10% Fetal Bovine Serum (FBS) and 1% penicillin, streptomycin and l-glutamine (PSG). For heavy methyl SILAC labelling, K562 cells were cultured for 2 weeks in RPMI-1640 with no methionine (Thermo Fisher Scientific, #A1451701) supplemented with 10% dialyzed FBS (Thermo Fisher Scientific, #A3382001), 1% PSG, and 0.1 mM of either light or heavy methionine (L-Methionine-(*methyl*-^13^C,d_3_), Sigma-Aldrich, #299154). Light and heavy methionine-labelled cells were mixed 1:1 and immediately prepared for Ribo Mega-SEC, as detailed below. A separate aliquot of heavy methionine-labelled cells was also prepared for Ribo Mega-SEC to verify heavy methyl incorporation.

### Ribo Mega-SEC

Ribosome populations were separated using Ribo Mega-SEC^59^, as described previously^60^, with some modifications. Briefly, 5 × 10^6^ K562 cells were incubated with 0.1 mg/mL cycloheximide for 5 min at 37 °C, rinsed in ice-cold PBS containing 0.1 mg/mL cycloheximide and resuspended in 250 μL Polysome Extraction Buffer (PEB, 20 mM HEPES-NaOH (pH 7.4), 130 mM NaCl, 10 mM MgCl_2_, 5% glycerol, 1% CHAPS (w/v), 0.2 mg/mL heparin, 2.5 mM DTT, 20 U/mL SUPERase in RNase inhibitor, 2 × cOmplete EDTA-free protease inhibitor, 0.1 mg/mL cycloheximide). For isolation of heavy methyl SILAC-labelled polysomes, cycloheximide was omitted from all steps. Cell were lysed by incubating on ice for 15 min with occasional vortexing, clarified by centrifugation at 17,000 *g* for 20 min, and clarified lysates were filtered through a 0.22 μm spin filter. Protein concentration was determined using a Pierce ™ 660 nm Protein Assay Kit (Thermo Scientific). Ribosome populations were then separated using size exclusion chromatography (SEC). Lysate containing ∼50 μg protein (polysome profiling) or ∼210 μg protein (hmSILAC polysome isolation) were separated on an Agilent Bio SEC-5 column (1000Å, 7.8 × 300 mm) for 45 min at 0.4 mL/min in SEC running buffer (20 mM HEPES-NaOH (pH 7.4), 60 mM NaCl, 10 mM MgCl_2_, 5% glycerol, 0.3% CHAPS (w/v), 2.5 mM DTT) using an Agilent 1260 Infinity Bio-Inert system. Absorbance at 254nm, 260nm and 280nm was measured throughout. For each sample, fractions were collected every 30 s from 15 min to 39 min, for a total of 48 fractions. Two repeat injections were carried out for hmSILAC samples. For quantifying polysomes and monosomes/60S, the area under the curve of the A_260_ chromatogram was measured to a zero baseline between 15 and 20.5 min (polysomes) or 20.5 to 23.5 min (monosomes/60S).

### Preparation of Ribo Mega-SEC fractions for mass spectrometry

For initial profiling of Ribo Mega-SEC fractions, 200 μL of fractions 1 to 40 (corresponding to 0.5 min fractions between 15 and 35 min) were individually prepared and digested with trypsin. For hmSILAC samples, Ribo Mega-SEC fractions corresponding to polysomes (fractions 1 to 11; 15 – 20.5 min) from both injections were pooled and 500 μL pooled polysomes was digested separately with each protease. For analyses of polysomes and monosomes/60S fractions from Cas9 only mock and *SMYD5* knockout cells, fractions corresponding to polysomes (15 to 20.5 min) or monosomes/60S (20.5 to 23.5 min) were pooled and digested with trypsin. For all samples, fractions (pooled or individual) were buffer exchanged, using Amicon Ultra-0.5 Centrifugal Filter Units with Ultracel-3 membrane, into either 50 mM NH_4_HCO_3_ (trypsin), 20 mM NH_4_HCO_3_ (LysC and GluC) or 100 mM Tris (pH 8.0)/10 mM CaCl_2_ (chymotrypsin). Samples were then reduced with 10 mM DTT for 30 min at 37 °C, alkylated with 15 mM chloroacetamide or iodoacetamide for 30 min at room temperature, and digested with 300 ng Sequencing Grade Modified Trypsin (Promega), 300 ng Glu-C (Promega), 300 ng rLys-C (Promega) or 600 ng Chymotrypsin (Promega) for 16-18 h at 37 °C (trypsin, GluC and LysC) or 25 °C (chymotrypsin). Peptides were then desalted with 50 mg tC18 Sep-Pak columns (Waters) and resuspended in 0.1% formic acid. Peptides were analysed on an Orbitrap Fusion Lumos Tribrid mass spectrometer (Thermo Scientific) as described below.

### Cloning, mutagenesis, expression and purification of SMYD5

The coding region of SMYD5 with a C-terminal hexahistidine tag, obtained as a gBlock (Integrated DNA Technologies), was cloned into pET15b by Gibson Assembly for bacterial expression. Point mutations of SMYD5 were generated using mutagenic primers, as described previously^61^, and SMYD5 deletions were generated by site-directed ligase-independent mutagenesis (SLIM)^62^ (see Table S5). Plasmids were transformed into Rosetta (DE3) *E. coli*, which were grown in lysogeny broth (LB) at 37 °C to an OD_600_ of ∼0.6, before expression of SMYD5 was induced with 1 mM IPTG and cells were grown for a further 5 h at 25 °C. Cells were harvested by centrifiguation and cell pellets kept at −80 °C until purification.

Wild-type or mutant SMYD5 was purified with either 1 mL Ni-NTA Superflow Cartridges (Qiagen) or His Mag Sepharose Ni resin (Cytiva), according to previous methods^63^. Briefly, *E. coli* cells were resuspended in His-tag purification buffer (50 mM sodium phosphate, 500 mM NaCl, 40 mM imidazole, 20% glycerol, pH 8), lysed by sonication (3 rounds of 40% amplitude for 30s, alternating 0.5 s on/0.5 s off) and clarified by centrifugation (21,000 *g* for 40 min at 4 °C). Clarified lysates were applied to equilibrated Ni-NTA cartridges or His Mag resin, which were washed with His-tag purification buffer before SMYD5 was eluted with His-tag elution buffer (50 mM sodium phosphate, 500 mM NaCl, 500 mM imidazole, pH 7.4). Purified SMYD5 was then buffer-exchanged into 50 mM sodium phosphate/200 mM NaCl (pH 7.4), and glycerol was added to 40% for storage at −80 °C.

### In vitro *methylation assays*

Full-length RPL40 (I-I-E-P-S-L-R-Q-L-A-Q-K-Y-N-C-D-K-M-I-C-R-K-C-Y-A-R-L-H-P-R-A-V-N-C-R-K-K-K-C-G-H-T-N-N-L-R-P-K-K-K-V-K) was synthesised using a MilliGen 9050 automated peptide synthesizer as described previously^64^.

For whole protein assays, synthetic RPL40 (5 μM), recombinant H3 (5 μM) or recombinant H4 (5 μM) were incubated with or without K562 lysate, Ribo Mega-SEC fractions, or recombinant WT or mutant SMYD5 (0.5 μM) in RPL40 *in vitro* methylation buffer (30 mM Tris-Cl pH 8.5, 80 mM MgCl_2_, 40 mM KCl, 1 mM EDTA, 10 mM DTT), in the presence of 50 μM D_3_-methyl-AdoMet ((RS)-S-Adenosyl-L-methionine-d3 tetra(p-toluenesulfonate) salt, Medical Isotopes) for 2 h at 37 °C. Reactions were stopped by addition of 6 × SDS loading buffer (350 mM Tris-Cl, 30% (v/v) glycerol, 10% (w/v) SDS, 600 mM DTT, 0.012% (w/v) bromophenol blue) and boiling for 10 min. Proteins were separated by SDS-PAGE, gels stained and visualised according to previous methods^63^. Gel bands corresponding to RPL40, H3 or H4 were then excised and destained in 25 mM NH_4_HCO_3_ / 50% acetonitrile, reduced with 10 mM DTT for 30-60 min at 37 °C and alkylated with 15 mM IAA for 30-60 min at room temperature. In cases where gel bands were subject to propionylation, this was done according to previous methods^65^. Finally, proteins were digested with 100 ng of trypsin for 18 h at 37 °C, before peptides were extracted from gel bands as described previously^3^.

For peptide assays, synthetic peptides (2 μM, ChinaPeptides) were incubated with or without recombinant SYMD5 (0.3 μM) in RPL40 *in vitro* methylation buffer, in the presence of 50 μM D_3_-methyl-AdoMet, for 1.5 h at 37 °C. Assays with H3K36 and H4K20 peptides were treated at 37 °C for a further 30 min, while RPL40K22 peptide assays were treated with 15 mM IAA for 30 min at room temperature. For assays with peptide mixtures for MT-MAMS, 500 μM D_3_-methyl-AdoMet was used instead, and the final concentration of peptide mixtures was 40 μM, resulting in 2 μM per peptide for all 20 substituted peptides. Assays were stopped by the addition of heptafluorobutyric acid to 1%, and peptides were desalted using 100 μL Bond Elut OMIX C18 tips (Agilent) according to the manufacturer’s instructions. Peptides were analysed by LC-MS/MS on an LTQ Orbitrap Velos (Thermo Scientific), according to previous methods^3^.

### *Generation of* SMYD5 *knockout cell lines*

To knockout *SMYD5*, 2 guide RNAs (gRNAs, Table S5) were designed using the Synthego CRISPR Design Tool (https://design.synthego.com/). Guide 1 was designed to target exon 2 and guide 2 was designed to target exon 3. The guides were ordered as Alt-R™ CRISPR-Cas9 crRNAs (IDT) and mixed with Alt-R CRISPR-Cas9 tracrRNA (IDT, #1072534) in equimolar concentrations to form the gRNA duplex. The gRNA duplex was then combined with the Alt-R™ S.p. Cas9-GFP V3 nuclease (IDT, #10008161) to form the RNP complex.

5 × 10^5^ K562 cells were transfected with the RNP complex using the Neon Transfection System (Thermo Fisher Scientific, #MPK5000). Cells were than cultured for 24 hours in RPMI-1640 supplemented with 10% FBS before fluorescence activated cell sorting. Live GFP positive cells were single-cell sorted into 96-well plates and left to grow (in RPMI-1640 with FBS and PSG) for 2 weeks or until colonies could be seen by eye. The genomic DNA for clonal populations of cells were then subjected to PCR (primers in Table S5) of the targeted regions (either exon 2 or exon 3) after DNA extraction using QuickExtract DNA Extraction Solution. PCR products were sent for Sanger sequencing followed by sequence alignment in SnapGene (https://www.snapgene.com/) to identify clones that resulted in frameshifts and SMYD5 protein truncations.

### *Analysis of RPL40 K22 methylation in* SMYD5 *knockouts*

Approximately 2-3 × 10^6^ cells of Cas9 only mock control or *SMYD5* KO K562 clones were pelleted and washed once with PBS before being resuspended in ice-cold RIPA buffer (150 mM NaCl, 1% Triton X-100, 0.5% Sodium Deoxycholate, 0.1% SDS, 50 mM Tris-HCl, pH 8) supplemented with 5 mM DTT, 1 mM PMSF, 10 ug/ml Leupeptin, and 10 ug/ml Aprotinin. Cells were agitated for 30 minutes at 4°C followed by centrifugation and collection of supernatants. Approximately 20 μg of whole cell lysate was separated by SDS-PAGE and the band corresponding to RPL40 was excised, subject to propionylation and digested with trypsin, as described above for *in vitro* assays. The methylation of RPL40 K22 was then monitored by parallel reaction monitory (PRM), as described below.

### Puromycylation assays

One million cells were seeded into a 6-well plate, before 10 μg/mL of puromycin was added and cells were incubated for 15 min at 37 °C. Cells were immediately placed in ice-water, washed with ice-cold PBS and lysed in RIPA buffer with protease inhibitors and 5 mM DTT, as above. Lysates were immunoblotted with an anti-puromycin antibody (MABE343, Merck, 1:2000 dilution in PBS-T with 5% (w/v) BSA) overnight at 4 °C, before incubation with rabbit anti-mouse IgG HRP antibody (ab97046, Abcam, 1:50,000 dilution in PBS-T with 5% (w/v) BSA) and detection by chemiluminescence according to previous methods^66^.

### Mass spectrometry

Peptide samples were analysed on an Orbitrap Fusion Lumos, Q Exactive Plus or LTQ Orbitrap Velos mass spectrometer. For all mass spectrometric analyses, peptides were separated by nanoLC and ionised by electrospray, as described previously^67^. Analyses on the Q Exactive Plus and LTQ Orbitrap Velos were carried out according to previous methods^3^, except without inclusion lists. Individual Ribo-Mega SEC fractions and hmSILAC-labelled polysomes were analysed on the Orbitrap Fusion Lumos according to previous methods^63^, except that for hmSILAC-labelled polysomes, peptides were separated over a 122 min gradient according to previous methods^67^. Deep proteomic analyses of Ribo Mega-SEC fractions F29-F33 were carried out on the Orbitrap Fusion Lumos, using HCD and ion trap detection for fragment ion scans, according to previous methods^67^. Parallel reaction monitoring (PRM) was carried out on the Orbitrap Fusion Lumos according to previous methods^35^, except using HCD fragmentation or EThcD fragmentation for the N-terminal RPL23A peptide. See Table S2 for peptides used for analysing RPL40 K22 methylation.

### Data analysis

Parallel reaction monitoring (PRM) and other targeted mass spectrometric data were analysed in Skyline (v. 21.2.0.568)^68^. For other analyses, raw mass spectrometry data were searched using MSFragger (v. 4.1)^69^ in FragPipe (v. 22.0) against the UniProt reference human proteome (UP000005640, downloaded 2023-06-26). For quantification of ribosomal proteins in Ribo Mega-SEC fractions, the “LFQ-MBR” FragPipe workflow was used. For analysis of hmSILAC-labelled polysomes, the “Default” FragPipe workflow was used, with changes to the protease and variable modification settings. Specifically, for trypsin samples: enzyme = stricttrypsin (enzyme cut = KR), allowed missed cleavages = 3; for LysC samples: enzyme = LysC (cut at K, no-cut P), allowed missed cleavages = 3; for GluC samples: enzyme cut = DE, allowed missed cleavages = 5; for chymotrypsin samples: enzyme cut = FYWLKRM, no-cut = P, allowed missed cleavages = 5. For variable modifications, mono-, di- and tri-methylation on lysine, arginine, histidine and the N-terminus were added, allowing up to 3 of each per peptide, i.e. 14.01565 - KRH[^, 28.0313 - KR[^ and 42.0469 - K[^. Due to the increased search space for GluC and chymotrypsin samples, methylation of R and H were searched separately to K and N-terminal methylation for these samples; i.e. variable modifications were 14.01565 – RH and 28.0313 – R, or separately, 14.01565 - K[^, 28.0313 - K[^ and 42.0469 - K[^. In combining search results for K/N-term and R/H methylation, the best scoring PSM was kept for each spectrum. Note that for determining sequence coverage of ribosomal proteins (Figure S2), methylation was not included as a variable modification. Putative methylpeptide PSMs were then analysed using MethylQuant (v. 1.0.0)^23^, to confirm the presence of hmSILAC pairs, according to previous methods^67^. Methylpeptide PSMs were then filtered for those with MethylQuant score ≥ 30 and a hmSILAC H/L ratio ≥ 0.5.

