## Supplementary material for "SMYD5 is a ribosomal methyltransferase which trimethylates RPL40 lysine 22 through recognition of a KXY motif": Document S1. Figures S1-S17 and Table S2.

**Table S2. Peptides used for detection and quantification of RPL40 K22 methylation by mass spectrometry.**

| **Peptide sequence** | **Modifications** | **Charge** | **Monoisotopic m/z** | **Purpose** |
|---|---|---|---|---|
| YNCDKMICRK | 2 × Carbamidomethyl (C), Oxidation (M) | 2+ | 702.3151 | Detecting unmethylated RPL40 K22 in samples not subject to propionylation. |
| KCYAR | Propionyl (K), Carbamidomethyl (C) | 2+ | 377.1892 | Detecting unmethylated RPL40 K22 in samples subject to propionylation. |
| KCYAR | $CD_3$-monomethyl (K), Carbamidomethyl (C) | 2+ | 357.6934 | Detecting monomethylated RPL40 K22 in *in vitro* assays not subject to propionylation. |
| KCYAR | $CD_3$-monomethyl + propionyl (K), Carbamidomethyl (C) | 2+ | 385.7065 | Detecting monomethylated RPL40 K22 in *in vitro* assays subject to propionylation. |
| KCYAR | $CD_3$-dimethyl (K), Carbamidomethyl (C) | 2+ | 366.2106 | Detecting dimethylated RPL40 K22 in *in vitro* assays (regardless of propionylation). |
| KCYAR | $CD_3$-trimethyl (K), Carbamidomethyl (C) | 2+ | 374.7279 | Detecting trimethylated RPL40 K22 in *in vitro* assays (regardless of propionylation). |
| KCYAR | Monomethyl + propionyl (K), Carbamidomethyl (C) | 2+ | 384.1971 | Detecting monomethylated RPL40 K22 from WT and *SMYD5* KO cells. |
| KCYAR | Dimethyl (K), Carbamidomethyl (C) | 2+ | 363.1918 | Detecting dimethylated RPL40 K22 |

| | | | | |
|---|---|---|---|---|
| | | | | from WT and *SMYD5* KO cells. |
| KCYAR | Trimethyl (K), Carbamidomethyl (C) | 2+ | 370.1996 | Detecting trimethylated RPL40 K22 from WT and *SMYD5* KO cells; also light RPL40 trimethylation from hmSILAC sample. |
| KCYAR | $^{13}CD_3$-trimethyl (K), Carbamidomethyl (C) | 2+ | 376.2329 | Detecting heavy-labelled RPL40 trimethylation from hmSILAC sample. |

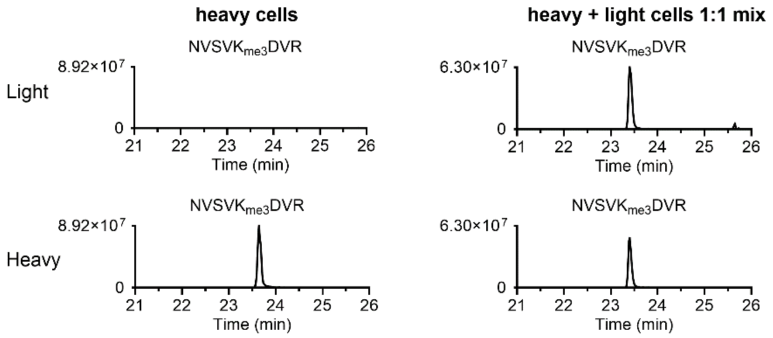


**Figure S1. Confirmation of heavy methyl SILAC labelling in K562 cells.**

Shown are extracted ion chromatograms (XICs) of light and heavy methylpeptides (sequence NVSVKDVR) corresponding to eEF1A K318 trimethylation, from purely heavy-labelled cells (left) or a 1:1 mix of light- and heavy-labelled cells (right).

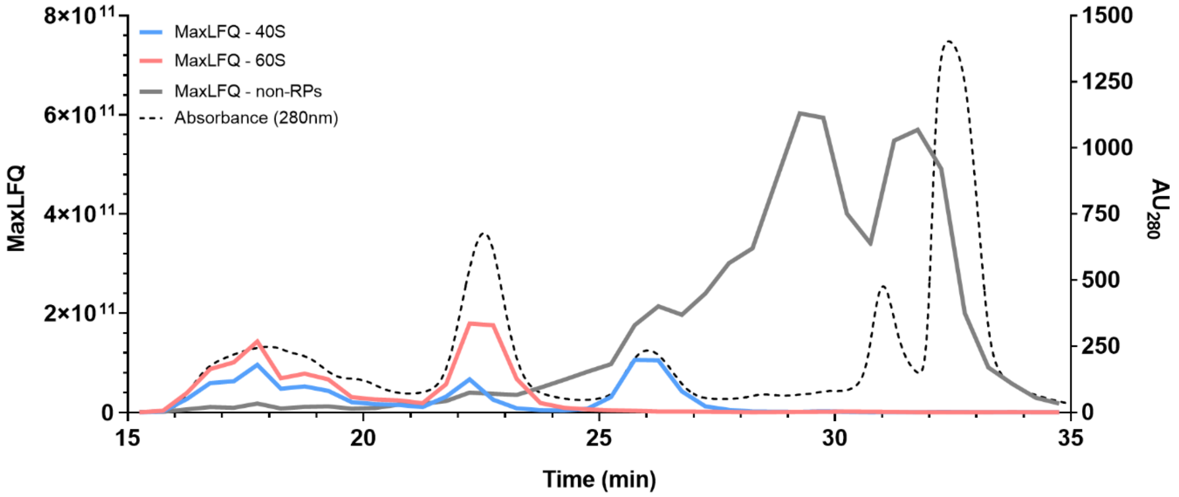


**Figure S2. Ribo Mega-SEC of human ribosomes using a 1000Å SEC column.**

Lysate containing ~250 µg of protein was separated by Ribo Mega-SEC at 0.4 mL/min on a BioSec-5 1000Å column. Fractions were collected every 0.5 min (i.e. 200 µL fractions) from 15 to 39 min. The 40 fractions corresponding to minutes 15 through 35 were digested with trypsin and analysed by LC-MS/MS. Summed MaxLFQ values for proteins of the large ribosomal subunit (60S), small ribosomal subunit (40S) or all other proteins (non-RPs) are shown in red, blue and grey respectively. The chromatogram of absorbance at 260 nm is shown as a dotted line.

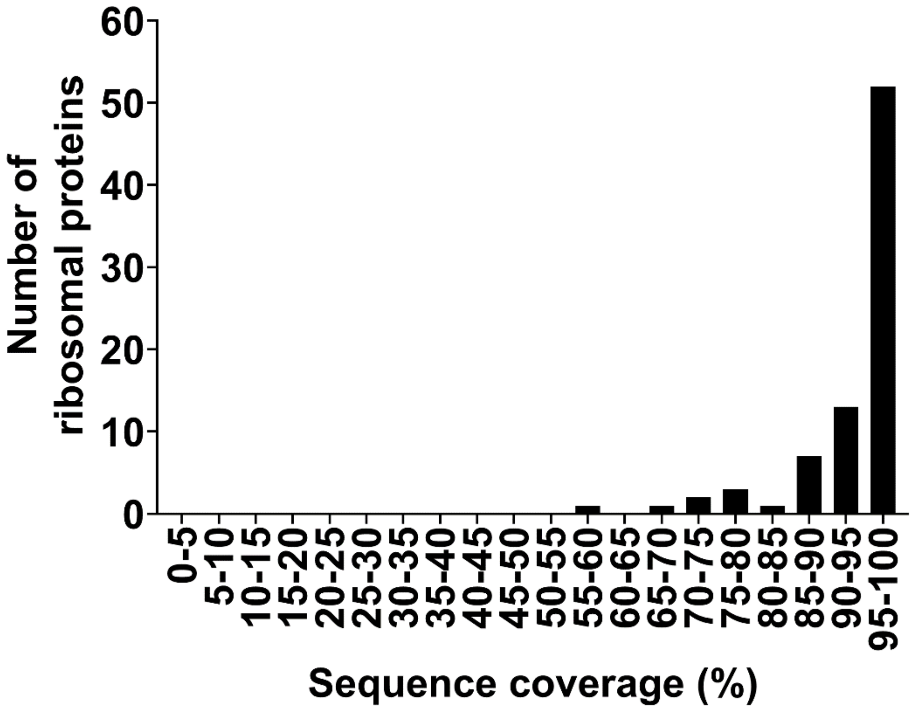


**Figure S3. High sequence coverage of ribosomal proteins.**

Histogram showing the sequence coverage of ribosomal proteins in the mass spectrometric analyses of hmSILAC labelled polysomes in Figure 1. Note that database searches for determining sequence coverage were carried out without methylation as a variable modification, to avoid false-positive methylpeptides contributing to the sequence coverage.

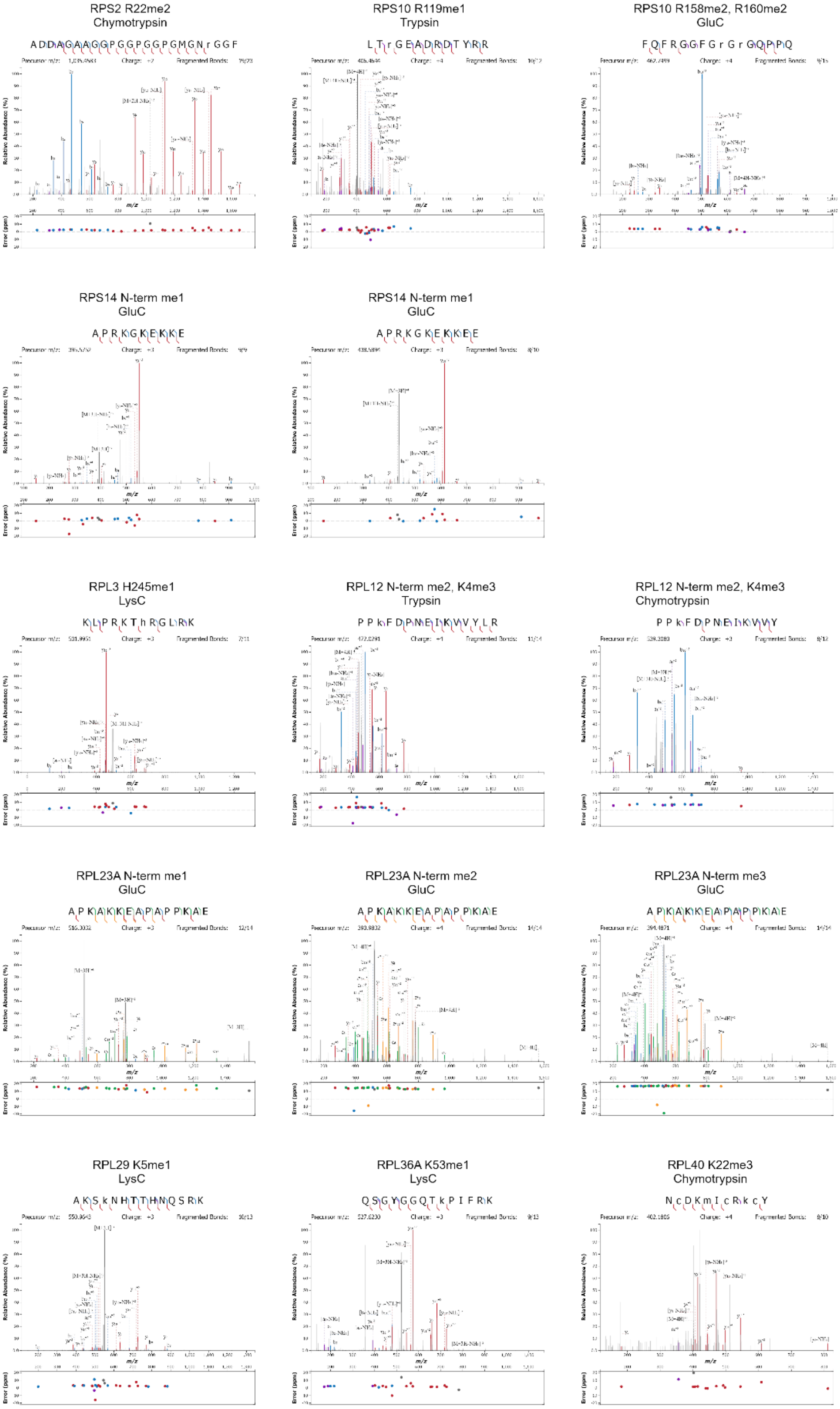


**Figure S4. Representative MS/MS spectra identifying ribosomal methylation sites.**

Chromatograms showing presence of light and heavy hmSILAC pairs are shown in Figure S5.

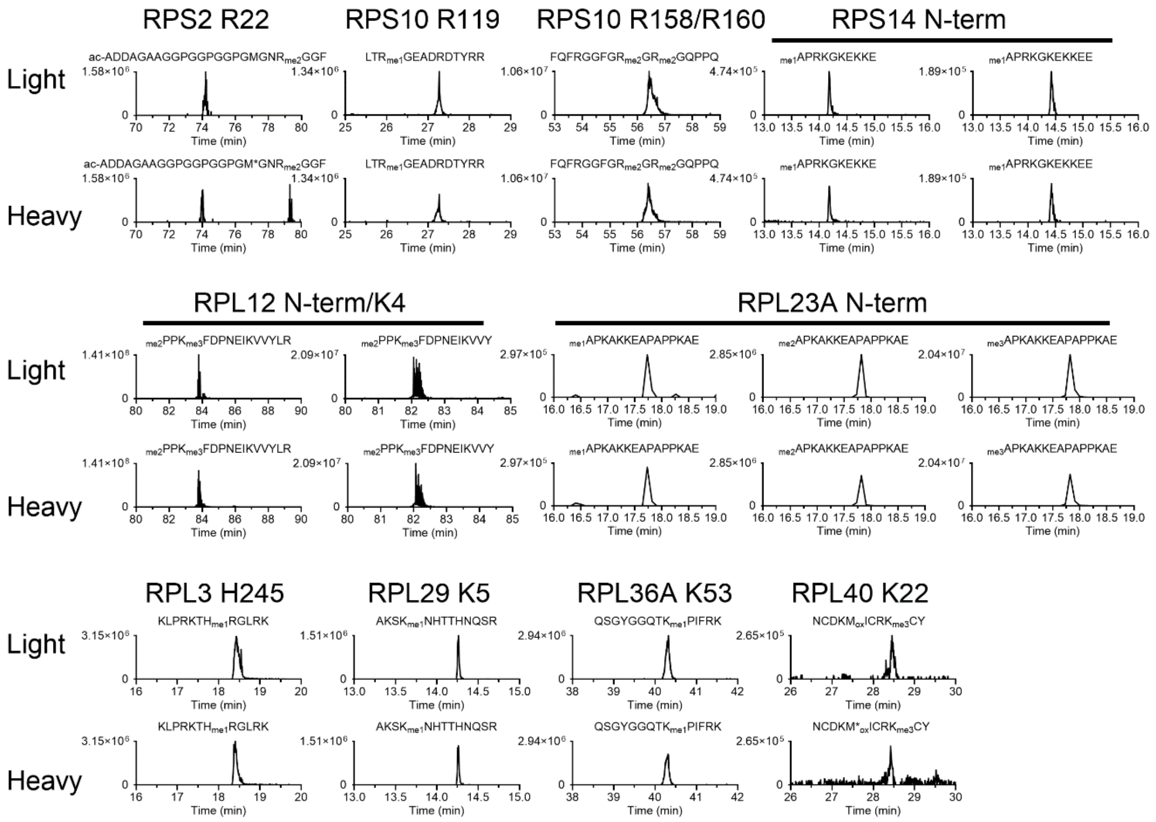


**Figure S5. Chromatograms of hmSILAC pairs for confirmed ribosomal protein methylation sites.**

Extraction ion chromatograms (XICs) of light and heavy methylpeptides were generated by taking 10 ppm windows around the m/z of each peptide in its light and heavy hmSILAC form. Asterisk (*) indicates the presence of a heavy-labelled methionine (+4 Da). MS/MS spectra for the light versions of all peptides are shown in Figure S4.

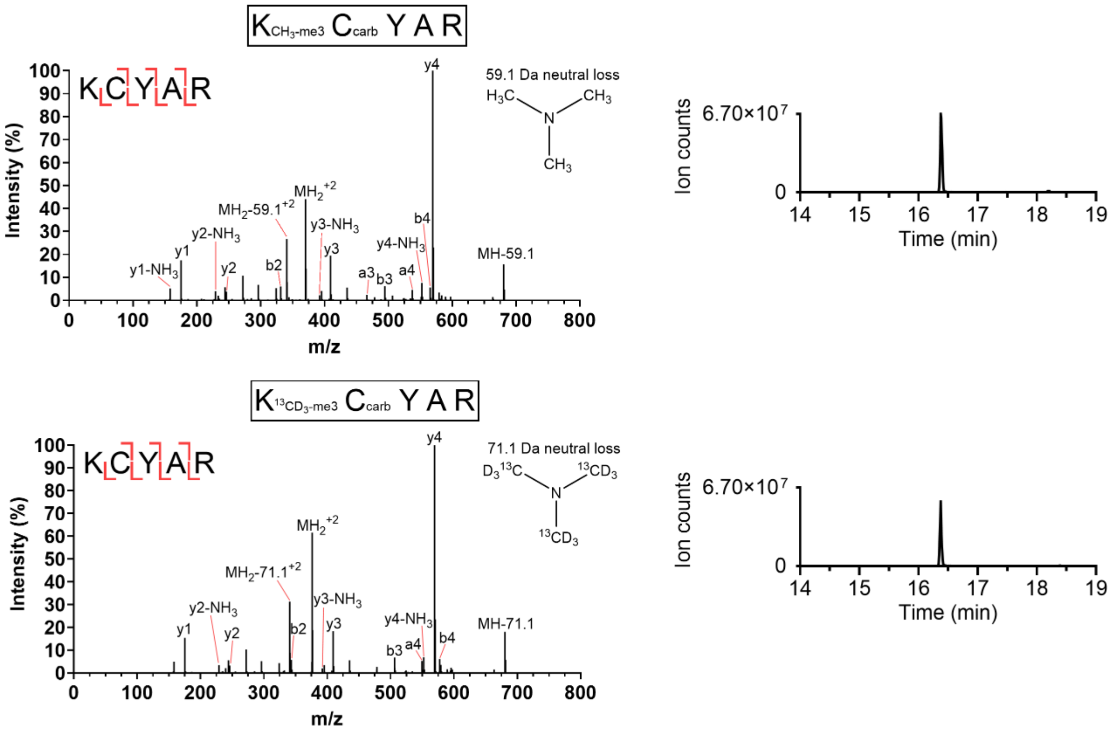


**Figure S6. Confirmation that RPL40 K22 is trimethylated in K562 polysomes.**

Shown on the left are Orbitrap MS/MS spectra of light (top) and heavy ($^{13}CD_3$-methyl) (bottom) trimethylated peptides containing RPL40 K22 (sequence KCYAR). Shown on the right are the corresponding XICs of these peptides, showing identical retention times and highly similar abundances.

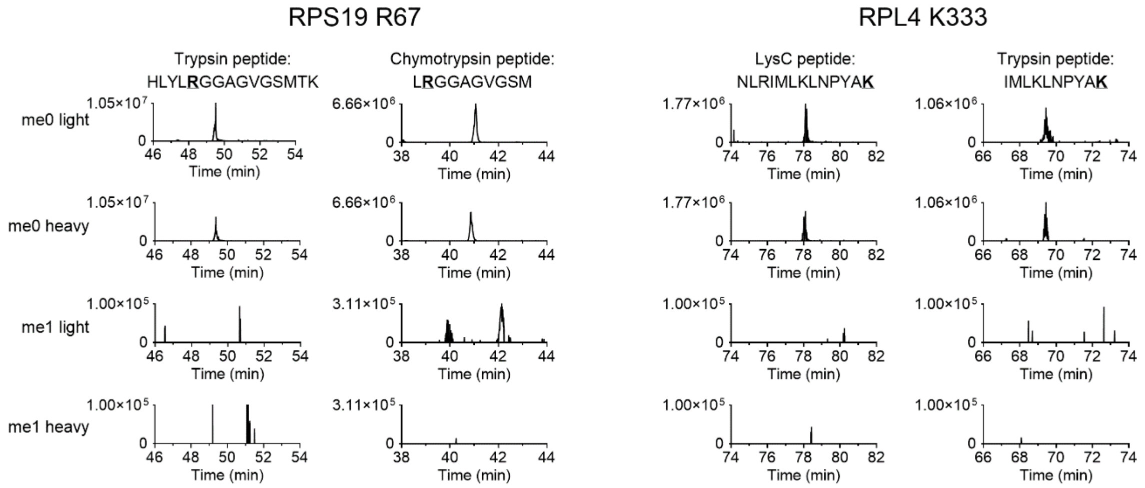


**Figure S7. RPS19 R67 and RPL4 K333 are not methylated in K562 polysomes.**

XICs of unmethylated light, unmethylated heavy, monomethylated light and dimethylated heavy peptides corresponding to putative methylation sites, showing an absence of any methylated peptides. Note both peptides have a heavy unmethylated version due to the presence of a methionine.

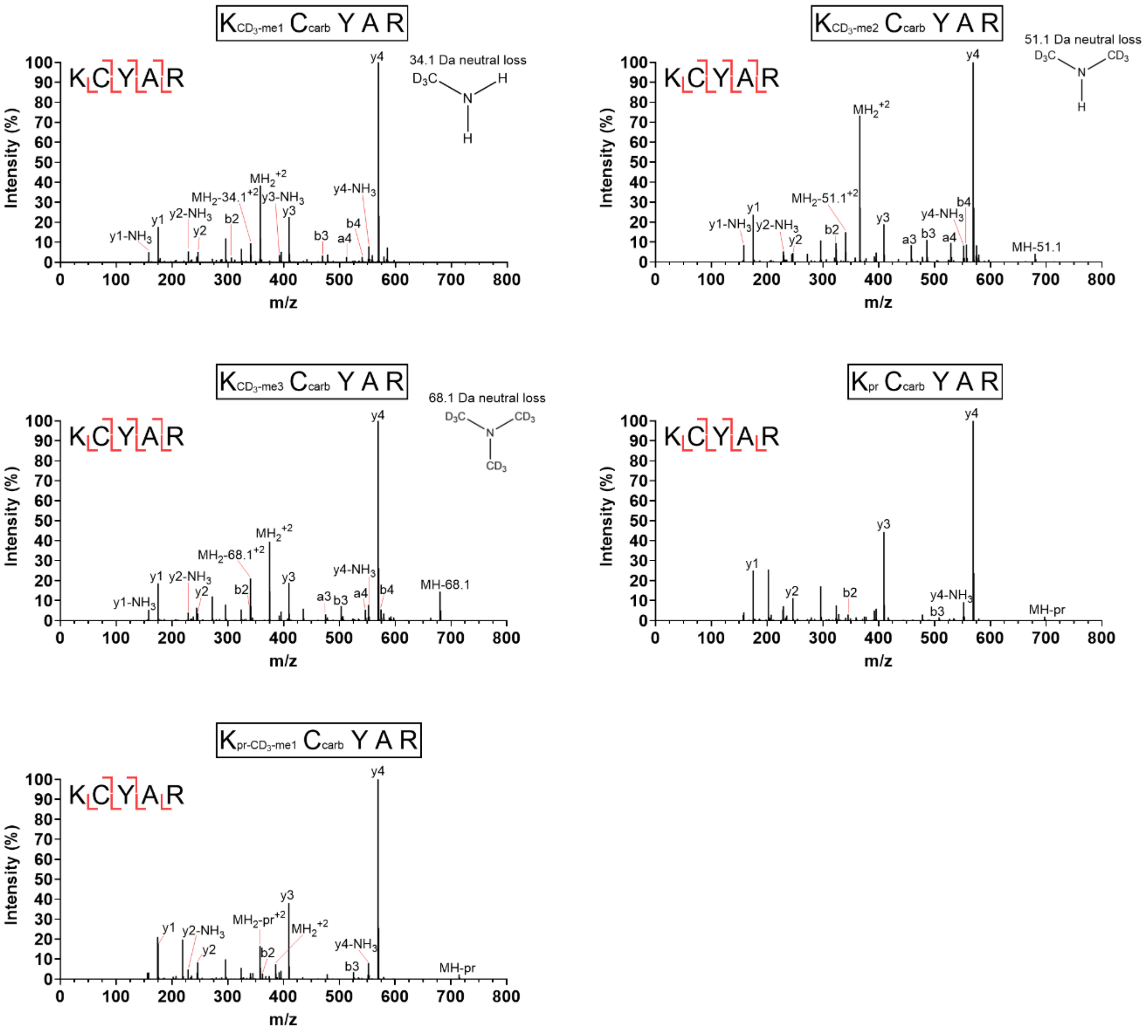


**Figure S8. Spectra of RPL40 K22-containing tryptic peptide KCYAR in all its D3-methylated and propionylated forms.**

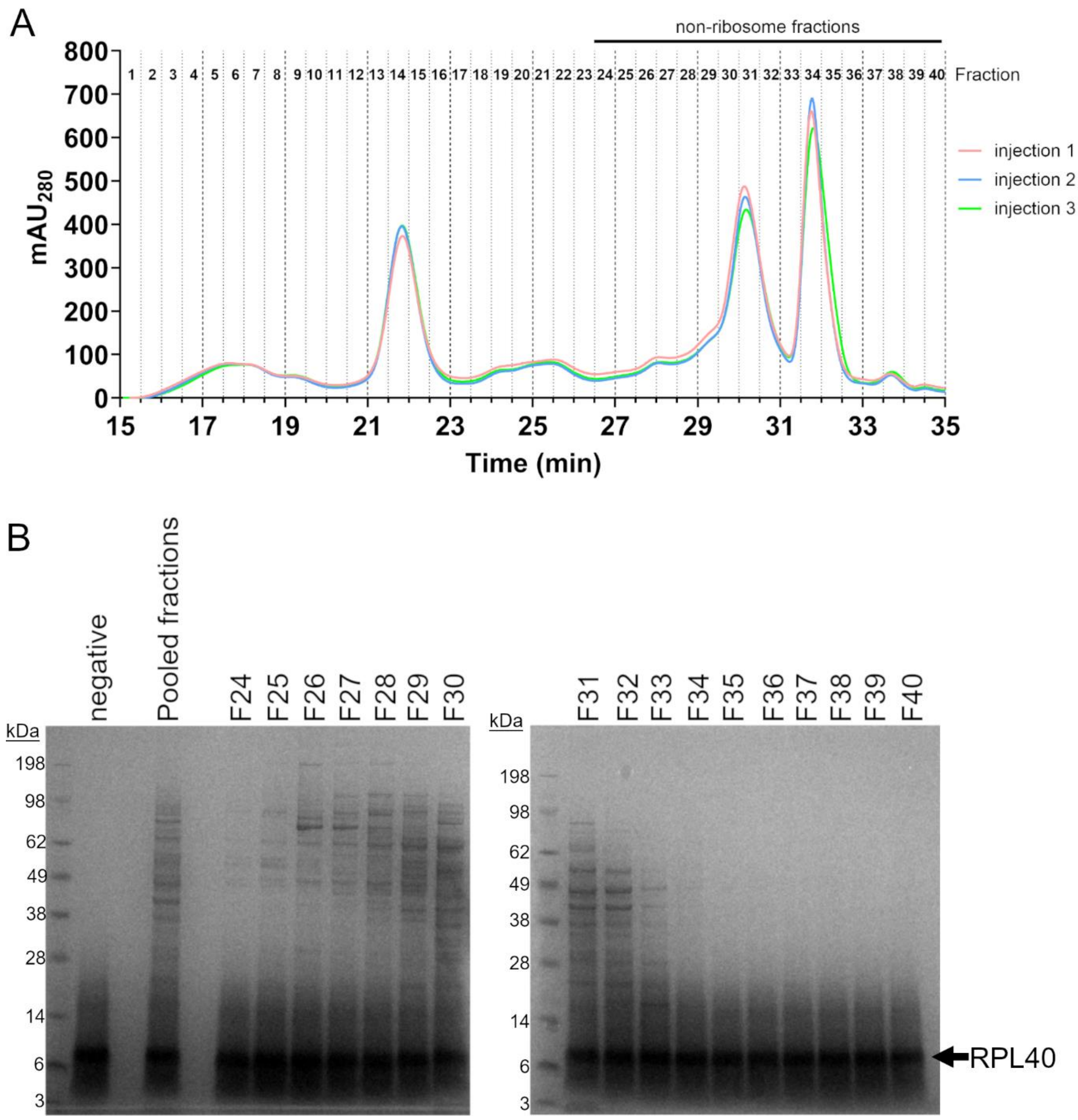


**Figure S9. *In vitro* methylation assays of RPL40 with non-ribosome-containing fractions from K562 lysate.**

**A)** Ribo Mega-SEC of K562 cells (unlabeled) was injected in triplicate runs. Shown are the chromatograms from all 3 runs. **B)** Non-ribosomal fractions (F24 to F40) were pooled from all 3 injections and concentrated. An equivalent volume of each fraction was incubated with synthetic RPL40 in the presence of $D_3$-methyl-AdoMet for 2h at 37 °C. Proteins were separated by SDS-PAGE, the band corresponding to RPL40 was excised and digested with trypsin, and RPL40 K22 methylation levels were analysed by LC-MS/MS.

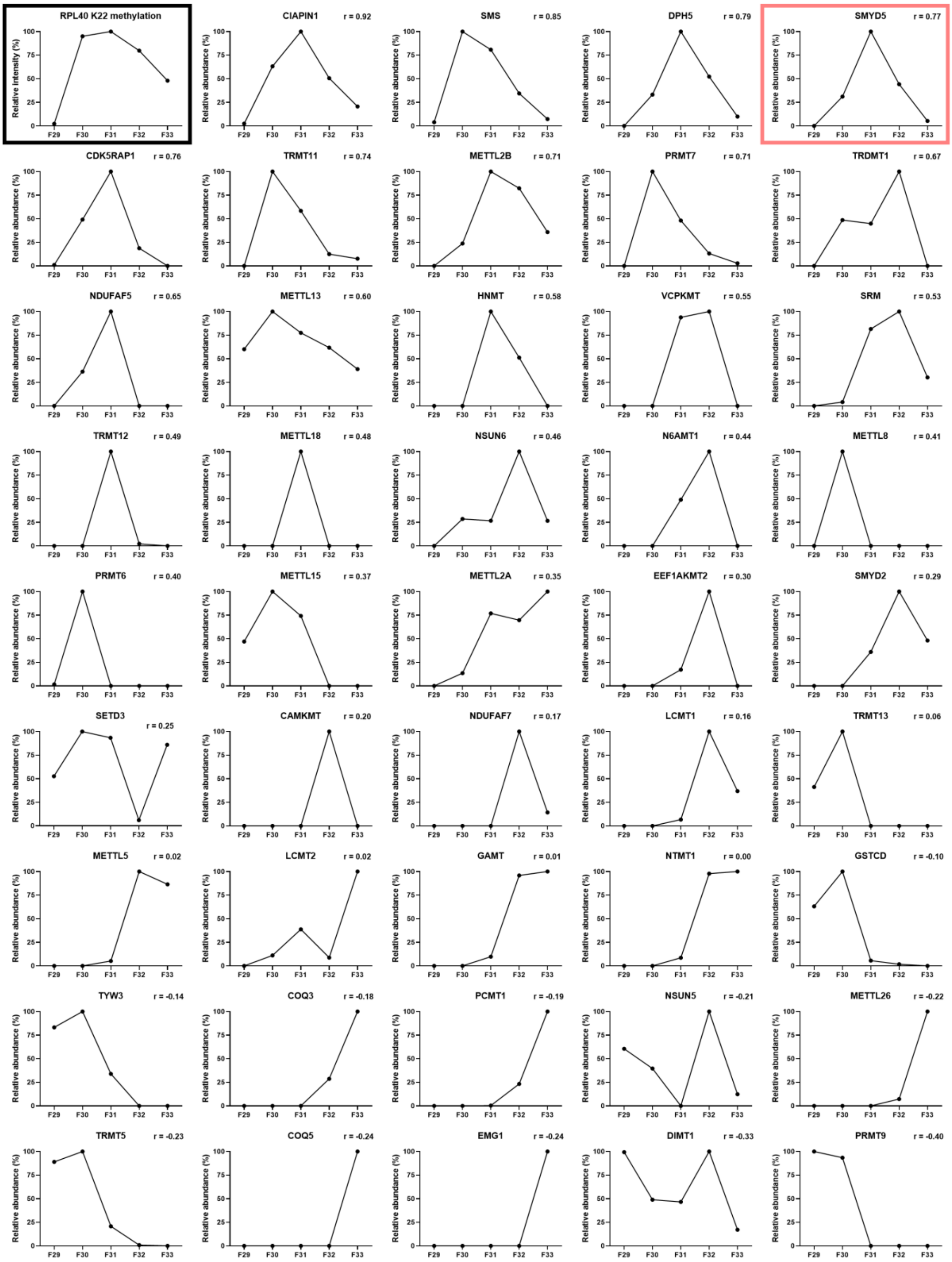


**Figure S10. Abundances of top 44 methyltransferase-like proteins correlating with RPL40 K22 methylation in RMSEC fractions F29-F33.**

Relative levels of RPL40 K22 methylation is shown in a black box in the top left. Methyltransferase abundance profiles are shown in order of decreasing correlation with RPL40 K22 methylation. The abundance profile of SMYD5 is indicated by a red box.

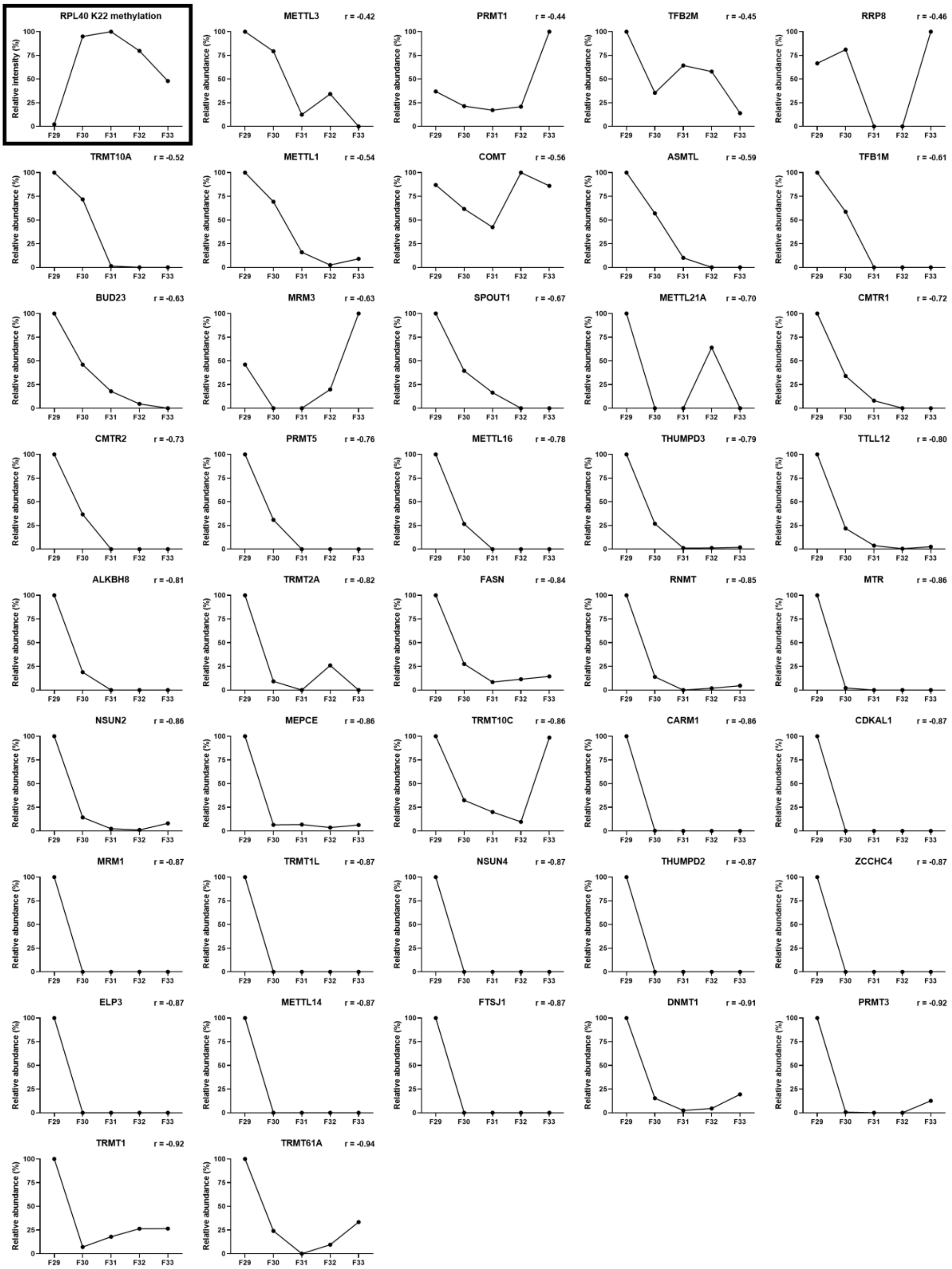


**Figure S11. Abundances of bottom 41 methyltransferase-like proteins correlating with RPL40 K22 methylation in RMSEC fractions F29-F33.**

Relative levels of RPL40 K22 methylation is shown in a black box in the top left. Methyltransferase abundance profiles are shown in order of decreasing correlation with RPL40 K22 methylation.

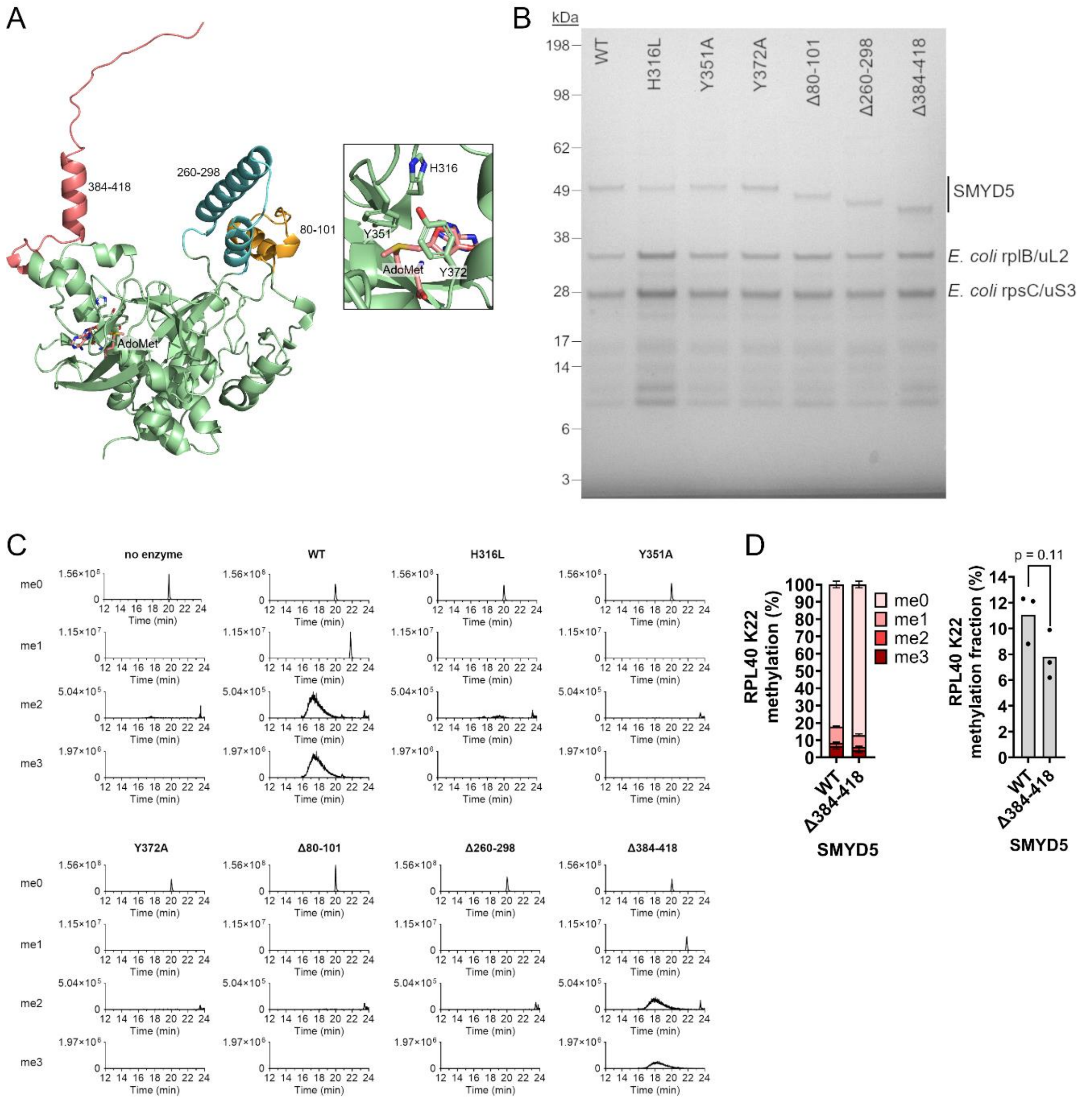


**Figure S12. Deletions and point mutations of SMYD5 affect its *in vitro* activity towards RPL40 K22.**

A) AlphaFold2 model of SMYD5 with AdoMet bound showing the regions deleted, including the M-insertion (80-101, orange), S-insertion (260-298, blue) and C-terminal poly-E acidic region (384-418, red). Inset: putative active site residues H316, Y351 and Y372. B) Purification of recombinant SMYD5 mutants. The two main co-purifying proteins from *E. coli* are shown. C) XICs of RPL40 K22-containing peptide KCYAR from *in vitro* methylation assays of RPL40 shown in Figure 2E. Peptide KCYAR, in either its unmethylated (me0), monomethylated (me1), dimethylated (me2) or trimethylated (me3) states was analysed by PRM. Only WT and Δ384-418 SMYD5 showed any activity towards RPL40 K22 *in vitro*. D) Triplicate methylation assays with WT or Δ384-418 SMYD5 against RPL40. Left: relative methylation levels of un-, mono-, di-, or tri-methylated RPL40 K22. Right: methylation fraction of RPL40 K22 relative to 100% trimethylation. A two-tailed t-test without equal variance was performed, showing a non-significant ($p = 0.14$) difference in methylation catalysed by WT or Δ384-418 SMYD5.

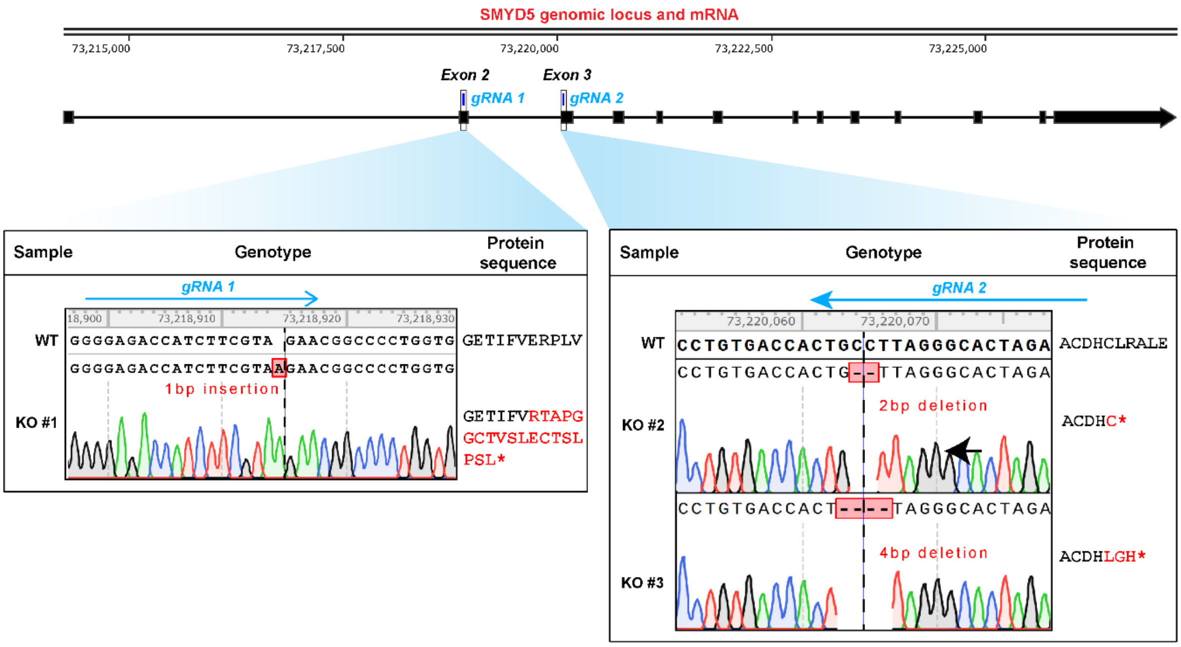


**Figure S13. *SMYD5* knockout in K562 cells.**

Schematic of the *SMYD5* genomic locus and mRNA with exons shown as black boxes. The two guide RNAs (gRNAs) are shown with a blue box above exons 2 and 3, respectively. K562 cells were transfected with Cas9 RNPs consisting of the Cas9 protein in complex with either gRNA 1 or gRNA 2. The genomic DNA from resulting clonal populations of K562 cells was then subjected to Sanger sequencing with primers spanning the targeted regions to look for successful knockouts. For each guide, a box is shown below which includes the WT sequence and aligned Sanger sequencing tracks, visualised in SnapGene, from the knockout clone(s) generated. A blue arrow above the WT sequence indicates the homology region for each gRNA with a black dotted line representing the Cas9 cut site. The predicted protein sequence starting from the region shown to a premature stop codon (*) is also listed on the right with red letters indicating altered amino acids or a premature stop codon.

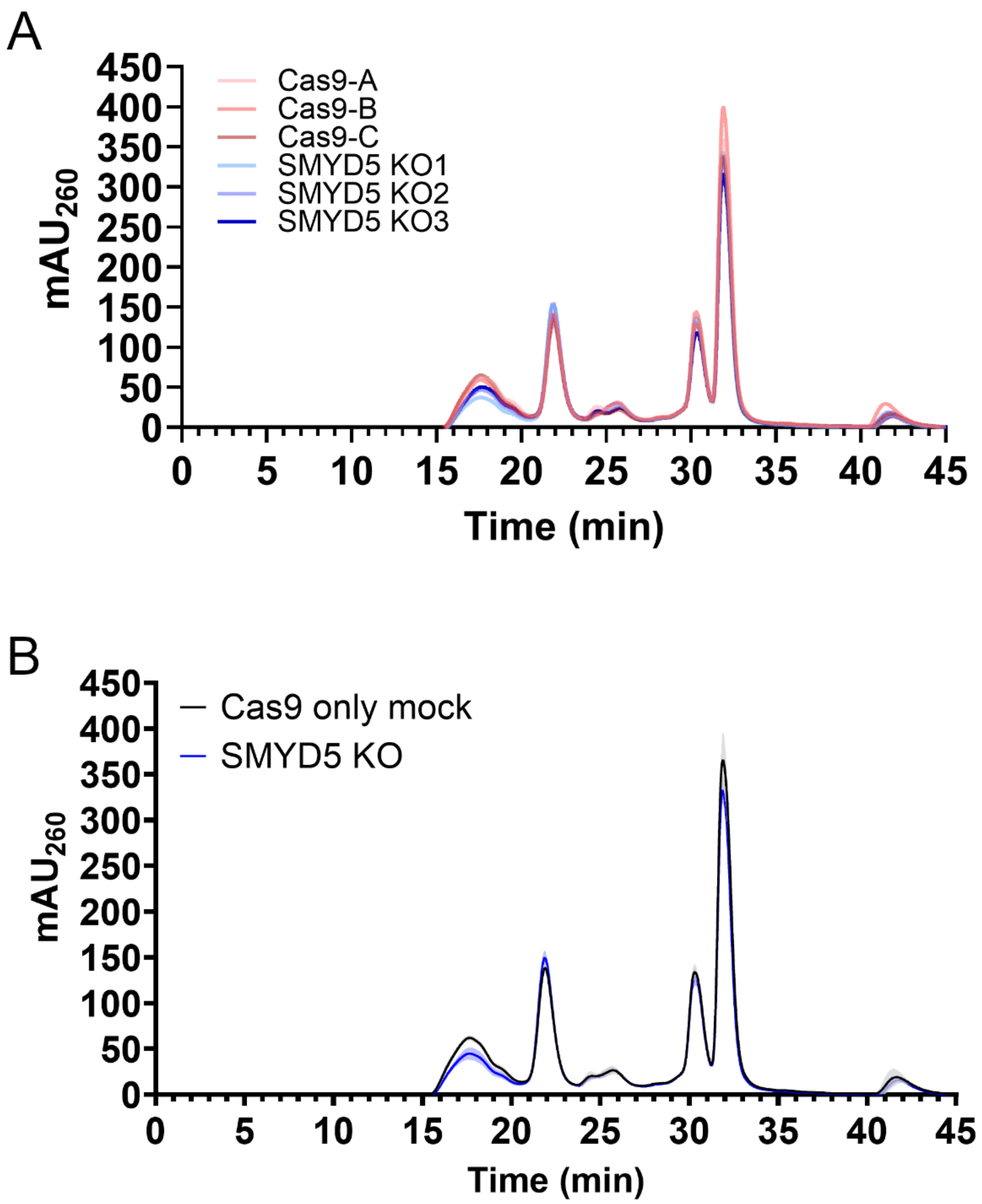


**Figure S14. Full Ribo Mega-SEC profiles for control and *SMYD5* knockout K562 cells.**

A) Individual Ribo Mega-SEC profiles for Cas9 only mock controls (Cas9-A, Cas9-B and Cas9-C) and *SMYD5* knockout cell lines (SMYD5 KO1, SMYD5 KO2 and SMYD5 KO3). B) Full averaged polysome Ribo Mega-SEC profiles for Cas9 only mock controls and *SMYD5* knockout cell lines, corresponding to Figure 3C. The shaded area represents one standard deviation (n = 3).

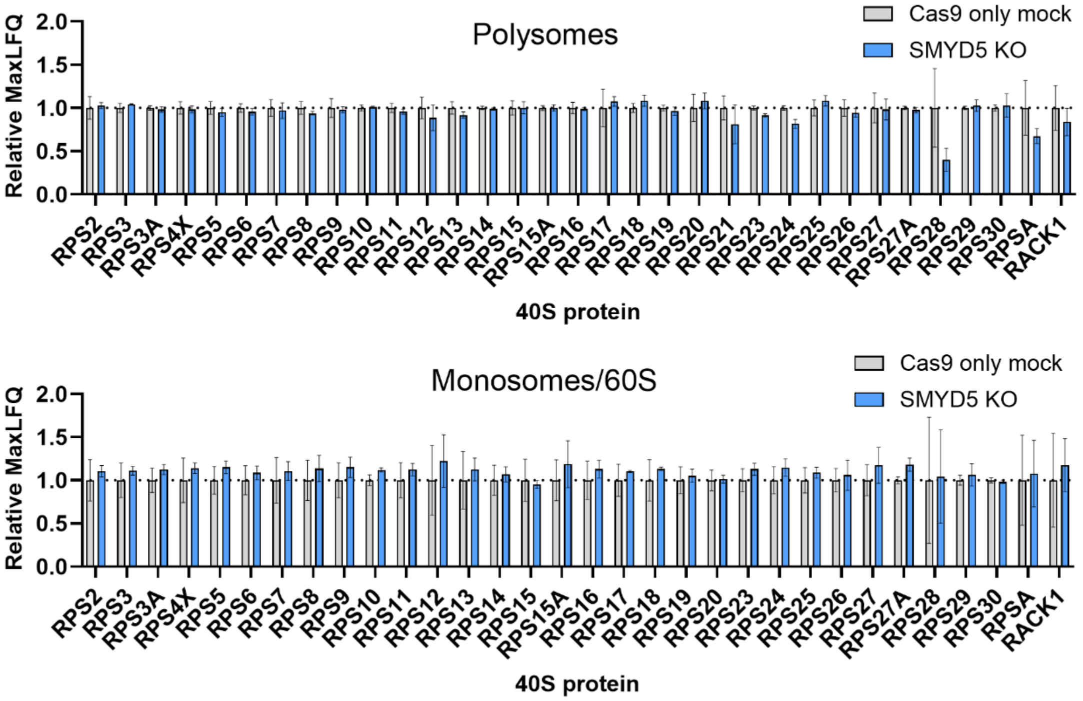


**Figure S15. Knockout of *SMYD5* does not affect 40S ribosomal protein levels in polysomes or monosomes/60S.**

Polysomes or monosomes/60S fractions from Ribo Mega-SEC analyses in Figure 3C were pooled and analysed by LC-MS/MS. MaxLFQ values for Cas9 only mock samples (n = 3) and *SMYD5* KO samples (n = 3) were normalised relative to the average MaxLFQ value for Cas9 only mock samples, separately for each ribosomal protein. Cas9 only mock and *SMYD5* KO samples were compared using unpaired t-tests with individual variances for each protein and multiple testing correction using the Holm-Šídák method. No significant differences were found ($p > 0.05$). Shown are 40S proteins; 60S proteins are shown in Figure 3E. Error bars show one SD.

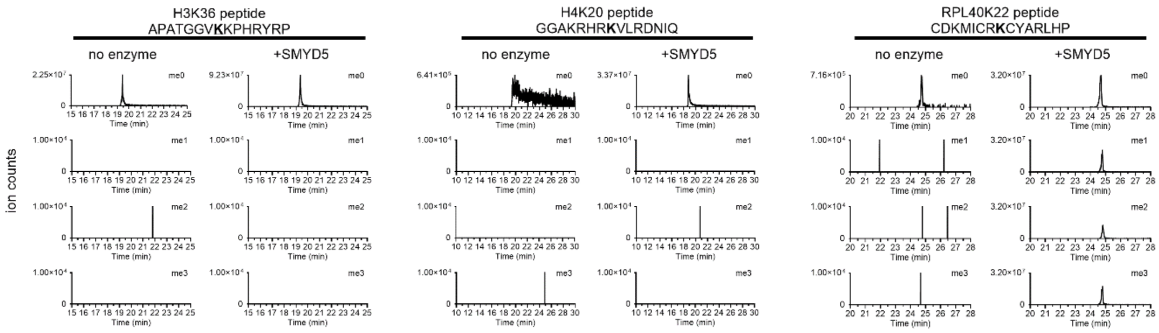


**Figure S16. Extracted ion chromatograms (XICs) from *in vitro* methylation assays of H3K36, H4K20 and RPL40K22 peptides with SMYD5.**

XICs were generated by taking windows (±10 ppm) around the predicted *m/z* of each peptide ion in its unmethylated (me0), monomethylated (me1), dimethylated (me2) or trimethylated (me3) state, wherein each methyl group adds 17.0345 Da. For the RPL40K22 peptide, carbamidomethylation of all three cysteines was also included. The triply-charged state was used for the H3K36 peptide and the quadruply-charged state for both the H4K20 and RPL40K22 peptides.

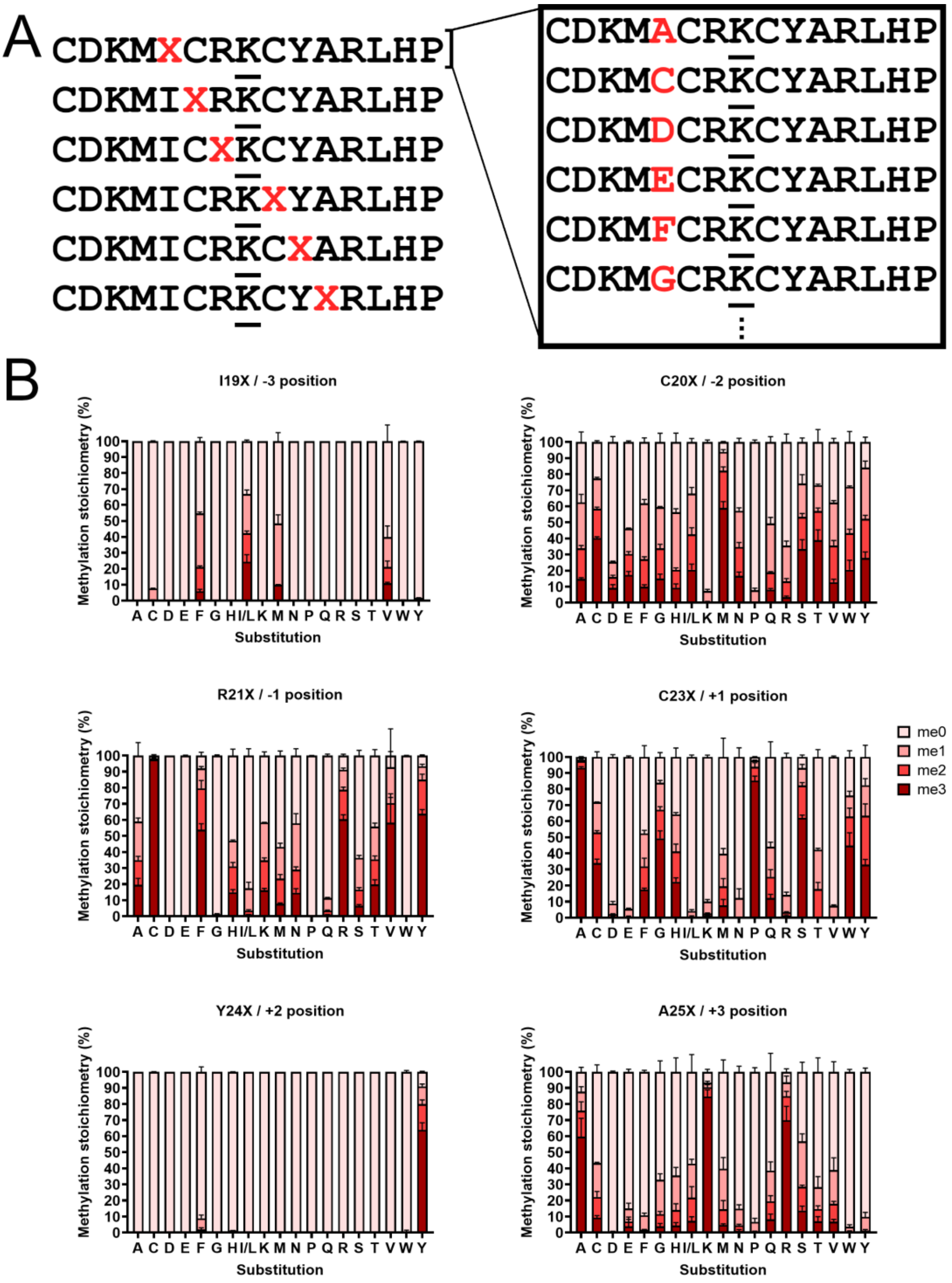


**Figure S17. MethylTransferase Motif Analysis by Mass Spectrometry (MT-MAMS) of SMYD5.**

A) Peptide mixtures used for MT-MAMS of SMYD5. Separately for each of the six positions around RPL40 K22 (shown as underlined), peptide mixtures were synthesised wherein the variable position (shown as a red X) is substituted to all 20 amino acids. Each peptide mixture was then assayed with SMYD5 and levels of methylation were detected by mass spectrometry. $D_3$-methyl-AdoMet was used for all assays to ensure the mass shift of methylation is unique and distinguishable from all amino acid substitutions. B) Methylation stoichiometries of all substituted peptides as catalysed by SMYD5. Error bars show one SD.
